# Augmin-mediated amplification of long-lived spindle microtubules directs plus-ends to kinetochores

**DOI:** 10.1101/501445

**Authors:** Ana F. David, Philippe Roudot, Wesley R. Legant, Eric Betzig, Gaudenz Danuser, Daniel W. Gerlich

## Abstract

Dividing cells reorganize their microtubule cytoskeleton into a bipolar spindle, which moves one set of sister chromatids to each nascent daughter cell. Early spindle assembly models postulated that spindle-pole-derived microtubules search the cytoplasmic space until they randomly encounter a kinetochore to form a stable attachment. More recent work uncovered several additional, centrosome-independent microtubule generation pathways, but the contributions of each pathway to spindle assembly have remained unclear. Here, we combined live microscopy and mathematical modeling to show that most microtubules nucleate at non-centrosomal regions in dividing human cells. Using a live-cell probe that selectively labels aged microtubule lattices, we demonstrate that the distribution of growing microtubule plus-ends can be almost entirely explained by Augmin-dependent amplification of long-lived microtubules. By ultra-fast 3D lattice light-sheet microscopy, we observed that this mechanism results in a strong directional bias of microtubule growth towards individual kinetochores. Our systematic quantification of spindle dynamics reveals highly coordinated microtubule growth during kinetochore-fiber assembly.

## Introduction

In dividing cells, the spindle apparatus congresses chromosomes to the cell equator and subsequently moves sister chromatids to the poles, so that each daughter cell inherits a complete copy of the genome. Spindles start forming upon mitotic entry, when the interphase microtubule (MT) network converts into an antiparallel, bipolar array (Heald and Khodjakov, 2015; Petry, 2016; Prosser and Pelletier, 2017). Vertebrate spindles attach a “fiber” of ~20-40 MTs to a confined region on each replicated sister chromatid, termed the kinetochore (Rieder, 1981; McDonald et al., 1992; McEwen et al., 1997; Deluca and Musacchio, 2012; Nixon et al., 2015; Walczak et al., 2010). Each pair of sister kinetochore-fibers binds to opposing spindle poles, enabling faithful chromosome segregation (Cimini et al., 2001; Tanaka, 2010).

Early spindle assembly models postulated that each MT in a kinetochore-fiber nucleates at one of the centrosomes to individually grow and capture kinetochores upon random encounter (Kirschner and Mitchison, 1986). Although stochastic capture events were observed in live cells (Hayden et al., 1990; Rieder and Alexander, 1990), mathematical modeling and computational simulations suggested that the probability of centrosomal MTs contacting all kinetochores within the typical duration of mitosis is extremely low (Wollman et al., 2005). Indeed, many MTs are now known to nucleate at pole-distal regions of the spindle, which is expected to increase the probability of kinetochore capture. MTs can nucleate in cytoplasmic regions surrounding chromosomes (Gruss et al., 2001; Sampath et al., 2004; Maresca et al., 2009; Meunier and Vernos, 2016; Petry and Vale, 2015; Scrofani et al., 2015), directly at kinetochores (Khodjakov et al., 2003; Sikirzhytski et al., 2018), or on the outer walls of existing MTs via the Augmin complex (Goshima et al., 2007; 2008; Kamasaki et al., 2013; Lawo et al., 2009; Petry et al., 2013; 2011).

The relative contributions of these alternative MT generation pathways to spindle assembly appear to vary across species and cell types (Meunier and Vernos, 2016). Centrosomal nucleation is thought to be the main source of spindle MTs in most animal cells (Prosser and Pelletier, 2017). Indeed, a comprehensive study in *Drosophila* embryonic cells confirmed this is the default dominant pathway, despite the fact that all act synergistically to ensure robust assembly of a bipolar spindle in a variety of perturbation conditions (Hayward et al., 2014). In mammalian cells, all of these pathways coexist (Gruss et al., 2002; Kalab et al., 2006; Kamasaki et al., 2013; Tulu et al., 2006). Yet, the contribution of each pathway to spindle assembly remains unclear. Importantly, the extent to which multiple processes are integrated in unperturbed mitosis is unknown.

Acentrosomal MT nucleation is best characterized in cytoplasmic extracts of *Xenopus* eggs, where it has long been thought to play a pivotal role in spindle assembly (Carazo-Salas et al., 1999; Kalab et al., 1999). Quantitative studies of the spatial distribution of MT plus-ends in this system demonstrate that acentrosomal nucleation makes an outstanding contribution to the assembly of the *Xenopus* meiotic spindle, allowing it to span radial distances of 50-300 μm (Petry et al., 2011; Brugués et al., 2012; Ishihara et al., 2014; Decker et al., 2018). The distribution of MT plus-ends in somatic cells from mammals (PtK-1 cells, (Tirnauer et al., 2002)) and Drosophila (Mahoney et al., 2006) appear to be inconsistent with a model of predominantly centrosomal nucleation. These observations call into question the early models of spindle assembly, which proposed the centrosome as the dominant source of MTs, and highlight the need for a formal quantification across multiple systems.

Here, we used live-cell microscopy and mathematical modeling to study spindle assembly in human somatic cells. We found that MT plus-ends arriving at metaphase chromosomes almost entirely originate from non-centrosomal regions via Augmin-mediated amplification of long-lived MT lattices. This pathway establishes a strong directional MT growth bias towards individual kinetochores, which promotes rapid formation of bioriented kinetochore-fibers. Our systematic quantification provides an integrated view on assembly and steady-state maintenance of vertebrate spindles.

## Results and Discussion

### A large fraction of the MTs that reach metaphase chromosomes does not originate from spindle poles

Current models for MT nucleation pathways predict the generation of plus-ends in distinct regions of the mitotic spindle. Whereas the Augmin-dependent and chromosome-dependent pathways can be expected to generate MTs across the entire spindle body (Kamasaki et al., 2013; Oh et al., 2016), centrosomal nucleation produces radial MT arrays. Considering the spherical geometry of these arrays, the density of MT plus-ends growing from centrosomes should decrease rapidly as a function of the distance from spindle poles. Hence, to determine the relative contribution of centrosomal and non-centrosomal MTs to spindle assembly in human cells, we sought to quantify MT plus-end density at increasing distances from spindle poles.

To this end, we acquired confocal time-lapse movies of HeLa cells (Fig. 1 A, Movie S1) and hTERT-RPE1 cells (Fig. S1 A, Movie S2) stably expressing enhanced green fluorescent protein-tagged EB3 (EB3-EGFP), which selectively binds growing MT plus-ends (Dragestein et al., 2008; Stepanova et al., 2003), and CENP-A-mCherry, a reference marker for kinetochores. Further, we stained chromosomes with the live-cell DNA marker SiR-Hoechst (Lukinavičius et al., 2015). In metaphase cells, the intensity and spatial distribution of EB3-EGFP fluorescence remained practically constant throughout one-minute movies, as did the overall morphology of the spindle (Movies S1 and S2). We therefore assumed photobleaching to be negligible and the MT plus-end distribution to be in a steady-state. This allowed us to project the EB3-EGFP fluorescence from all time frames to generate a smooth map of the average MT plus-end density (Fig. 1 B; Fig. S1 A). We then quantified EB3-EGFP mean fluorescence in sectors extending from the spindle pole to the spindle center to determine MT plus end densities. Considering a 3D spherical geometry of MTs growing radially from centrosomes, the density of MT plus-ends is expected to decrease by the inverse of the squared distance from spindle poles. If all MTs radiating from the centrosome rim grow to a length of 5 μm, the density of plus-ends measured at that distance should thus fall to 1% (Fig. 1 C, black). The EB3-EGFP fluorescence measured in interpolar spindle regions, however, decreased less than 40% near the spindle pole and then remained roughly constant (Fig. 1 C, green). For distances between 1 and 5 μm from the pole, the mean fluorescence stabilized at 63.1 ± 9.9% (mean ± s.d.) in HeLa cells (Fig. 1 C) and 69.6 ± 9.3% in hTERT-RPE1 cells (Fig. S1 B). This indicates that only a small fraction of MT plus-ends arriving at the metaphase plate can be part of the radial MT network originating from centrosomes.

**Figure 1.**
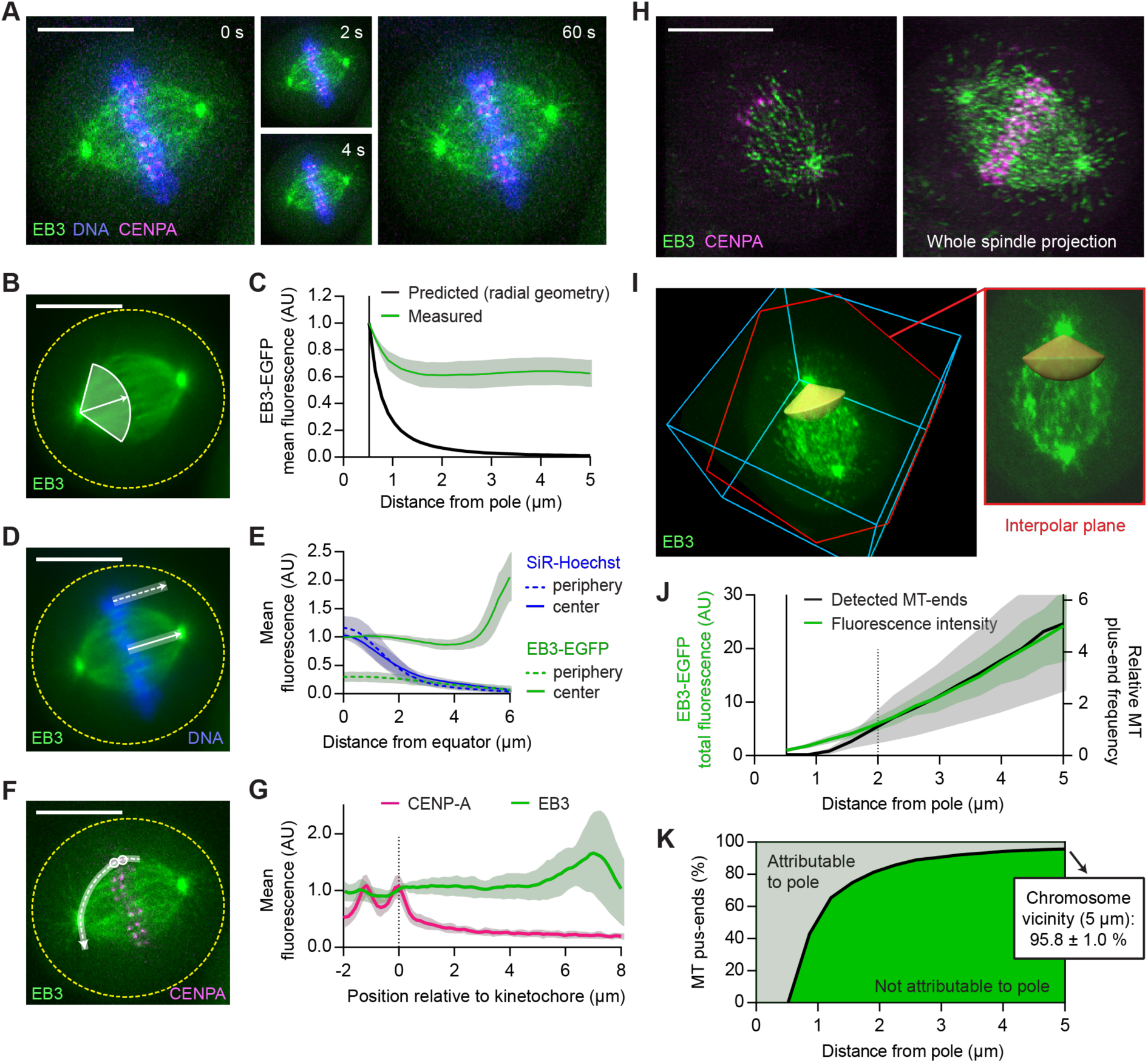
The large majority of MT plus-ends that reach metaphase chromosomes do not originate from spindle poles. (A-G) Live-cell confocal microscopy of HeLa cells expressing EB3-EGFP (green) and mCherry-CENPA (magenta), stained with SiR-Hoechst (blue). (A) Metaphase cell imaged for 1 min at 2 s/frame, see Movie S1. (B and C) Quantification of MT plus-end distribution from the spindle poles (see Materials and Methods for details). (B) All time frames were registered to correct for spindle rotation (see Movie S3) and projected to obtain mean-intensity images. EB3-EGFP fluorescence was measured in interpolar spindle regions, along a series of circumferential lines of increasing radius centered around the spindle poles. The quantification region extends from the pole to the spindle center (full white line, arrow indicates direction of pole-pole axis). (C) Mean EB3-EGFP fluorescence measured as in B (n = 25 cells; individual measurements normalized to centrosome rim). Black line indicates predicted signal dilution by radial geometry. (D and E) Quantification of EB3-EGFP and SiR-Hoechst fluorescence across the chromosome-cytoplasm boundary. (D) Mean intensities were profiled along lines placed either inside the spindle (full white line) or in its immediate periphery (dotted white line); results plotted in (E; n = 25 cells). (F) Quantification of fluorescence of EB3-EGFP and mCherry-CENPA along curved lines connecting pairs of sister-kinetochores (dashed white line) to one of the spindle poles, in maximum-intensity projections of 3 sequential movie frames. (G) Mean intensity profiles, aligned to the midpoint between sister-kinetochores (n = 42 profiles in 7 cells). (H-K) Hela cells imaged during metaphase by 3D lattice light-sheet microscopy (n = 11). (H) Maximum intensity projections of 5 (left) and 60 (right) consecutive slices of a deconvolved z-stack. (I) EB3-EGFP fluorescence was measured in non-deconvolved stacks, inside conical regions of interest defined around the interpolar axis (yellow). Shown is the same cell as in H; Bounding box: 23x24x11 μm. The slice highlighted in red follows the plane defined by the spindle poles and a random kinetochore. (J) Distribution of MT plus-ends in interpolar spindle regions, estimated from the EB3-EGFP fluorescence measured as in I (green) or from the EB3-EGFP particles detected as in Fig. S1 F (black). Fluorescence intensities were normalized to the centrosome rim; the count profiles shown in Fig. S1 G were normalized to 2 μm from the spindle poles. (K) Fractions of MT plus-ends attributable (light-green) and not attributable (dark-green) to nucleation at the spindle poles, as a function of distance from pole. Computed from the fluorescence measurements shown in J as detailed in Materials and Methods. Lines and shaded areas denote mean ± s.d., respectively. Scale bars, 10 μm. Yellow dotted lines indicate cell boundaries.

Given the dynamic instability of MT plus-ends (Mitchison and Kirschner, 1984), not all MTs nucleated at centrosomes are expected to reach metaphase chromosomes. Dynamic instability should therefore further decrease the fraction of interpolar MT plus-ends that can be attributed to centrosomal origin. A model taking both dynamic instability and spherical geometry into account predicts that 98.8 ± 0.2% (mean ± s.d.) of the MT plus-ends observed at 5 μm from the pole must originate from non-centrosomal regions (Fig. S1 C; see Materials and Methods for details). A high abundance of acentrosomal MT nucleation was previously shown to drive the formation of huge MT asters in egg extracts of *X. laevis* (Brugués et al., 2012; Ishihara et al., 2014; Decker et al., 2018). Our data show that also in metaphase spindles of much smaller human somatic cells most MTs are acentrosomal, fully compensating for the spherical-geometry-imposed dispersion of radial MT arrays.

To determine the predominant sites of non-centrosomal MT generation, we first investigated whether MT plus-ends might be locally enriched around the metaphase plate. We measured EB3-EGFP fluorescence across the chromatin-cytoplasm boundary (Fig. 1 D). Line profiles placed perpendicular to the metaphase plate showed only marginal enrichment in direct vicinity to SiR-Hoechst-stained chromosomes, irrespective of their position inside or in the periphery of the main spindle (Fig. 1 E and S1 D). Furthermore, analysis of EB3-EGFP fluorescence along curved line profiles connecting kinetochores with spindle poles also showed no substantial local increase when approaching kinetochores (Fig. 1 F-G and S1 E). Hence, MT plus ends are not locally enriched around chromosomes or kinetochores.

To further quantify the MT plus-end distributions, we aimed to visualize the entire 3D volume of metaphase spindles. Analyzing the distribution of EB3-EGFP in a 3D volume, rather than a thin optical section, should allow us to exclude the contribution of out-of-focus fluorescence to our MT plus-end density profiles. The photosensitivity of mitotic cells limits the analysis of MT plus-ends in 3D by conventional confocal microscopy. Lattice light-sheet microscopy offers high spatial resolution and ultra-fast image acquisition rates at minimal phototoxicity (Chen et al., 2014). Moreover, it uses scanned Bessel beams to illuminate extremely thin sample sections. This technology enabled us to visualize EB3-EGFP-labeled MT plus-ends across the entire volume of metaphase HeLa cells, along with our kinetochore marker mCherry-CENP-A (Fig 1 H).

We analyzed MT plus-end distributions in lattice light-sheet microscopy data by automatically detecting both spindle poles in each movie frame, using an in-house developed software (see Methods for details). We then quantified the EB3-EGFP fluorescence intensity in interpolar regions of the spindle, inside two pole-focused conical regions of interest (Fig. 1 I). The total fluorescence increased steadily as a function of distance from the pole (Fig. 1 J, green), such that 95.8 ± 1.0% (mean ± s.d.) of the signal sampled at 5 μm distance could not be attributed to centrosomal origin (Fig. 1 K). This result agrees with our previous estimates (Fig. S1 C), indicating that the contribution of out-of-focus fluorescence was small and confocal images can in principle be used to characterize the distribution of EB3-EGFP in the spindle.

To determine whether the measured fluorescence intensity profiles accurately reflect the distribution of MT plus-ends, we next aimed to detect individual EB3-EGFP-labeled particles in the spindle (Fig. S1, F and G). At pole-proximal regions, the density of MT plus-ends was too high for reliable particle detection. However, in pole-distal regions, where density was low enough for reliable particle detection, the observed distribution closely matched that of the total fluorescence measurements (Fig. 1 J, black). This indicates that the contribution of background fluorescence to these measurements is negligible, thus a quantification of fluorescence intensity can in principle be used to estimate the distribution of MT plus-ends in the metaphase sindle. Overall, these data indicate that centrosomes generate only a very minor fraction of the MTs whose plus-ends arrive at the spindle equator.

### Most MT plus-ends in metaphase spindles are generated in an Augmin-dependent manner

Given the continuous increase in MT plus-end number from spindle poles to the equator (Fig 1 G), we hypothesized that they might be predominantly generated by a MT-lattice-dependent MT generation mechanism. Two such mechanisms have been described: MT severing by Katanin (McNally and Vale, 1993; McNally et al., 2006; Roll-Mecak and McNally, 2010), and Augmin-mediated nucleation on the lattices of preexisting MTs. Transfection of siRNAs targeting Katanin (Dong et al., 2017) did not affect the distribution of EB3-EGFP fluorescence (Fig. S1, H and I). We hence considered Augmin-mediated nucleation as a major source of MT plus ends, as previously proposed (Goshima et al., 2007; 2008; Uehara et al., 2009; Kamasaki et al., 2013; Lawo et al., 2009; Petry et al., 2013; Zhu et al., 2008).

To quantify the contribution of the Augmin complex to MT plus-end generation, we depleted its HAUS6 subunit by RNAi (Fig. 2, A and B) and recorded 3D lattice light-sheet microscopy movies of metaphase HeLa cells 48 h after siRNA transfection (Fig. 2, C and D). Consistent with previous observations in fixed cells (Zhu et al., 2008), HAUS6-depletion noticeably reduced the amount of MT plus-ends in interpolar regions of the spindle, compared to control RNAi cells (Fig. 2 E). Depletion of HAUS6 reduced the accumulation of MT plus-ends from the pole by 2-fold. Importantly, the accumulation rate is substantially reduced along the entire length of the spindle (Fig. 2 F), indicating a widespread defect in MT plus-end generation within the spindle body. We observed a similar reduction in MT plus-end density in HAUS6-depleted hTERT-RPE1 cells (Fig S2, A and B).

**Figure 2.**
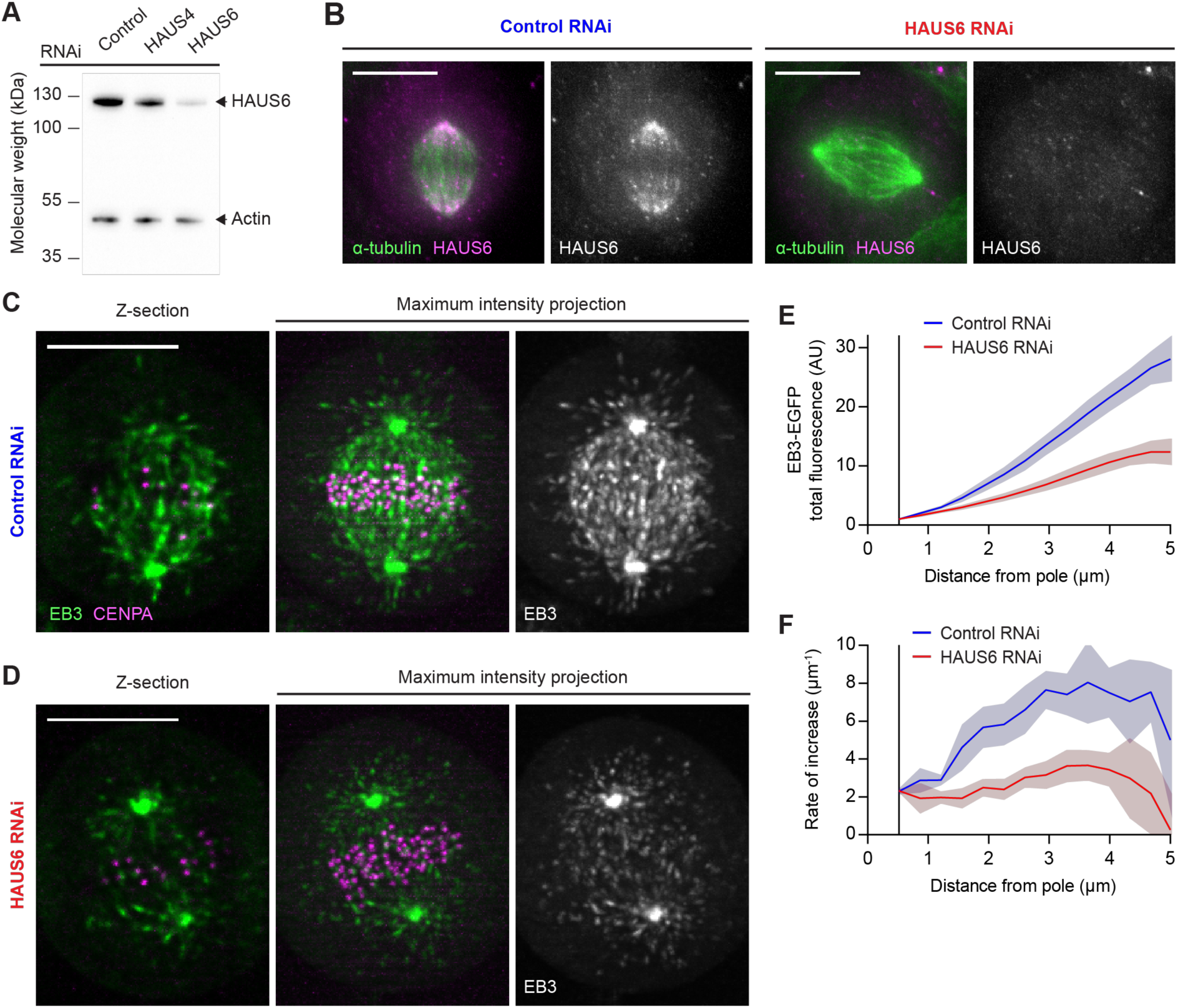
Most MT plus-ends in metaphase spindles are generated in an Augmin-dependent manner. (A) HAUS6 protein levels analyzed by Western Blotting, in HeLa cells transfected with control, HAUS4-targeting or HAUS6-targeting siRNAs. (B). HAUS6 and α-tubulin visualized by immunofluorescence in metaphase spindles of HeLa cells transfected with control or HAUS6-targeting siRNAs. (C-D) 3D Lattice light-sheet microscopy of HeLa cells expressing EB3-EGFP (green) and mCherry-CENPA (magenta), transfected with either (C) non-targeting Control siRNAs or (D) siRNAs targeting HAUS6. 2.5-minute movies of metaphase cells were acquired at 1 s/frame. Deconvolved images are shown. (E) EB3-EGFP fluorescence intensities measured in interpolar regions of the spindle as in Fig. 1 I, for control and HAUS6 RNAi cells (n = 7 and 11 cells, respectively, collected in 2 independent experiments). (F) First derivative ofEB3-EGFP fluorescence profiles shown in (E), indicating rate of increase for MT plus-end numbers. Lines and shaded areas denote mean ± s.d., respectively. Scale bars, 10 μm.

To test whether the reduced MT plus-end density was a specific effect of Augmin depletion, we examined cells transfected with siRNAs targeting HAUS4, another subunit of this complex. Metaphase spindles of HAUS4- or HAUS6-depleted cells showed similarly reduced MT plus-end density, compared with those of control RNAi cells (Fig. S2, C and D). When measured along kinetochore-to-pole trajectories (as in Fig. 1 F-G), MT plus-end density was reduced overall but showed no peak in the vicinity of kinetochores (Fig. S2 E). Thus, we could find no evidence of HAUS6-independent MT generation at kinetochores. To corroborate that the observed spindle perturbations were caused by on-target depletion of Augmin, we transfected cells with a plasmid encoding an siRNA-resistant mutant of HAUS6 and monitored its expression by visualization of EGFP fluorescence (Fig. S2 F; see Methods for details). These cells maintained normal spindles after transfection of HAUS6-targeting siRNAs, whereas control cells expressing only EGFP showed the expected decrease in MT plus-end density (Fig. S2 G). Hence, addback of the siRNA-resistant HAUS6 could completely suppress the RNAi phenotype. Together, our results indicate that Augmin-dependent nucleation is the main source of MT plus-ends in interpolar regions of the metaphase spindle.

Augmin is thought to amplify a preexisting MT network by generating new MTs on the lattice of others (Goshima et al., 2008; Petry et al., 2013). To investigate whether this might explain the MT plus-end distribution we observed in metaphase spindles, we first mapped MT lattice densities in HeLa cells stably expressing EGFP-α-tubulin (Fig. S3 A). The integrated fluorescence along circumferential line segments showed that the total number of MTs in interpolar regions increases steadily with the distance from the pole (Fig. S3 B). Thus, not all MTs of the metaphase spindle body extend to the poles, consistent with previous reports (Kamasaki et al., 2013; Nixon et al., 2017).

**Figure 3.**
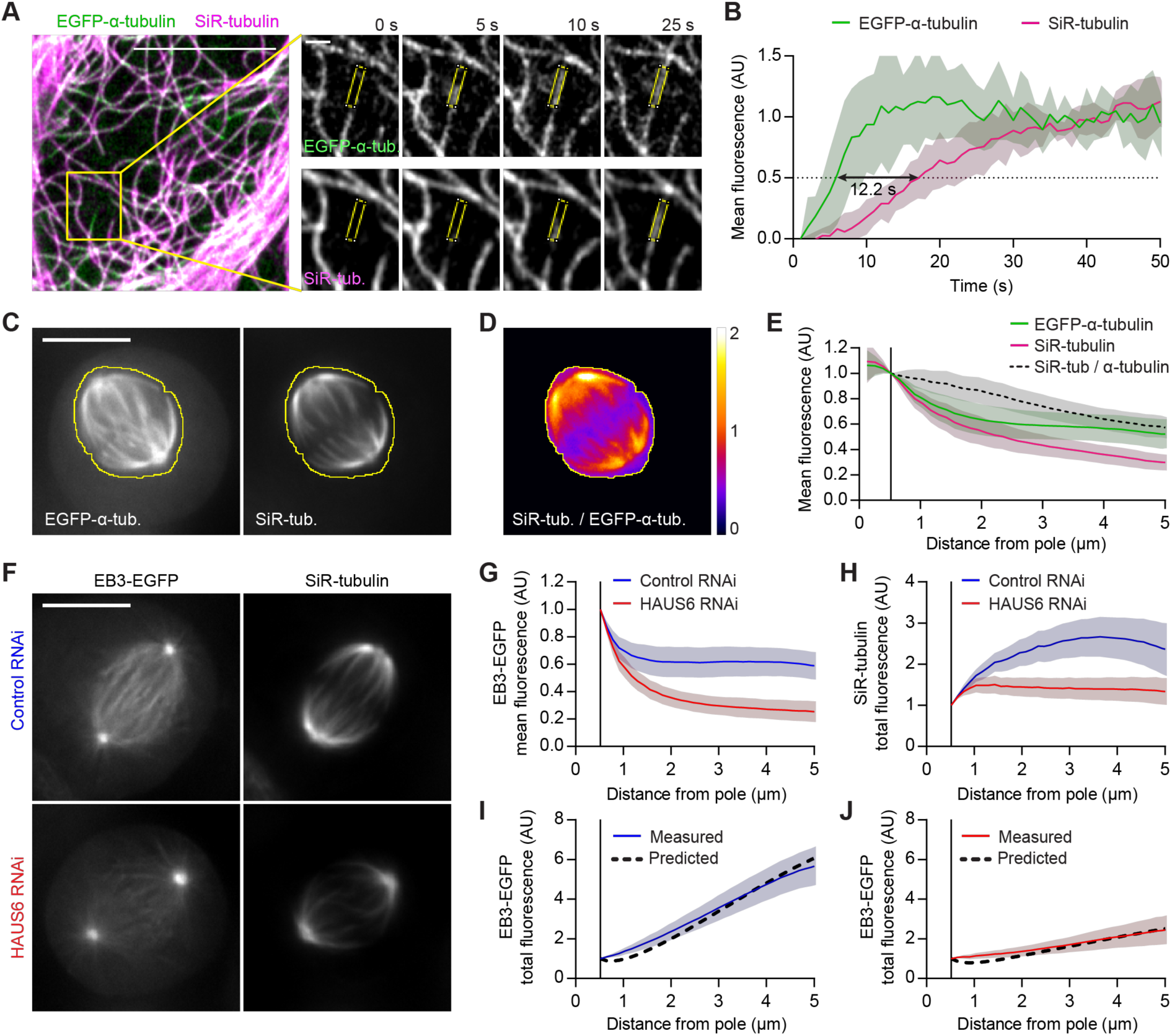
Uniform amplification of aged MTs by Augmin explains most of the plus-end distribution in the metaphase spindle. (A) Live interphase HeLa cell expressing EGFP-α-tubulin incubated with 100 nM SiR-tubulin imaged at 1 s/frame. Right panels show a typical MT growth event; mean fluorescence intensities were quantified in ROIs as illustrated (yellow). (B) Quantification of mean fluorescence in growing MTs as illustrated in (A); (n = 13 MTs in 9 cells; individual profiles normalized to the mean values measured at t> 30 s). Black arrow indicates lag time between the 50% fluorescence value of EGFP-α-tubulin and SiR-tubulin, respectively. (C-E) Live EGFP-α-tubulin-expressing HeLa cell incubated with 50 nM SiR-tubulin. Yellow line denotes spindle boundary. (D) Ratiometric image of SiR-tubulin / EGFP-α-tubulin, calculated based on temporal projections of 30 s movies for cell shown in (C). (E) Mean fluorescence intensities in interpolar regions quantified as in Fig. 1 B-C (n = 35 cells). (F) EB3-EGFP-expressing HeLa cells transfected with either non-targeting (control) or HAUS6-targeting siRNAs were incubated with 100 nM SiR-tubulin; average-intensity projections of registered movies. (G and H) EB3-EGFP (G) and SiR-tubulin (H) fluorescence, quantified as in Fig. 1 B-C (n = 32 and 35 cells for control and HAUS6 RNAi cells, respectively). SiR-tubulin fluorescence in interpolar spindle regions (H) served as amplification template in a mathematical simulation of MT plus-end density (see Figure S3 C, Materials and Methods for details). (I and J) Comparison between the predicted distributions of MT plus-ends (black dashed line) and the measured EB3-EGFP total fluorescence, for (I) control and (J) HAUS6 RNAi cells. Lines and shaded areas denote mean ± s.d., respectively. Scale bars, 10 μm unless otherwise indicated.

We implemented a mathematical model for MT plus-end distribution that adds the plus-ends expected to arise from amplification of a template MT network to the centrosome-generated plus-ends. The molecular factors driving the amplification were set to distribute evenly across all template lattices to promote local plus-end generation at a constant rate, such that the instantaneous frequency of pole-distal plus-end generation is a linear function of the MT template distribution (Fig. S3 C, see Methods). We used the measured distributions of EGFP-α-tubulin as template and predicted MT plus-end distributions in the interpolar regions of metaphase spindles. The simulations predicted a sharp increase in MT plus-end numbers along the spindle axis, partly explaining the distribution measured in EB3-EGFP-expressing HeLa cells (Fig. S3 D). However, the fit was not perfect, predicting a relative excess of MT plus-ends in pole-distal regions. Thus, although most MT plus-ends in the metaphase spindle are Augmin-dependent, their radial distribution does not fit well to a uniform amplification of all MT lattices.

### SiR-tubulin, a marker for aged MT lattices

Cold-induced depolymerization of unstable MTs in metaphase spindles does not substantially impact HAUS6 abundance in interpolar regions (Zhu et al., 2008), suggesting that Augmin predominantly localizes to stable MT lattices. Thus, we hypothesized that our model simulations would yield a more accurate fit if only long-lived MT lattices were used as a template for Augmin-mediated amplification. As cold-treatment perturbs the spindle geometry, we aimed to establish a live-cell marker for selective visualization of long-lived MT lattices.

*In vitro*, the fluorescent taxol derivative Flutax-1 binds to microtubules with relatively slow kinetics (Díaz et al., 2000). We thus hypothesized that the non-toxic, plasma membrane-permeant variant of Flutax-1, SiR-tubulin (Lukinavičius et al., 2014), might also bind to the lattice of MTs with a detectable delay and hence label MTs in an age-dependent manner. To test this, we incubated live HeLa cells expressing EGFP-α-tubulin in the presence of SiR-tubulin and recorded confocal time-lapse movies of the bottom surface of interphase cells to resolve individual MTs as they polymerize (Fig. 3 A and Movie S4). Quantification of the mean fluorescence along the lattice of growing MTs revealed that SiR-tubulin indeed labelled MT lattices with a delay of 12.2 ± 1.5 s (mean ± s.e.m.) relative to their polymerization, as visualized by EGFP-α-tubulin (Fig. 3 B). Because of this delay, SiR-tubulin did not label those stretches of MT lattice that persisted for less than 12 s before their disassembly owing to dynamic instability (Fig. S3, E and F). Hence, SiR-tubulin enables selective visualization of long-lived MT lattices.

To map the distribution of SiR-tubulin-stained MTs in metaphase spindles, we recorded one-minute time-lapse movies cells expressing EGFP-α-tubulin, as a reference marker for total MTs (Fig. 3 C). We found that central spindle regions contained substantially lower ratios of SiR-tubulin to EGFP-α-tubulin fluorescence, compared to pole-proximal regions (Fig. 3 D). The average fraction of stained lattices decreased steadily with increasing distance from the spindle poles, such that at 5 μm it is only 57 ± 8.3% of the fraction measured at the centrosome rim (Fig. 3 E). These observations suggest that aged MT lattices are enriched in pole-proximal spindle regions.

To corroborate this observation, we aimed to visualize aged MT lattices with an alternative probe. Acetylation is a posttranslational modification of α-tubulin thought to accumulate on MT lattices in an age-dependent manner (Szyk et al., 2014; Janke and Montagnac, 2017). We imaged metaphase spindles in fixed cells co-stained with an antibody targeting acetylated tubulin and another that recognizes α-tubulin irrespective of acetylation (Fig. S3 G). We found that pole-proximal spindle regions were substantially enriched in acetylated tubulin, whose relative distribution closely resembled that of SiR-tubulin (Fig. S3 H, compare with Fig. 3 E). This validates SiR-tubulin as a marker for aged MT lattices, and demonstrates that aged MT lattices accumulate in spindle pole-proximal regions.

### The MT plus-end distribution in metaphase spindles fits a uniform amplification of long-lived MTs

The relatively low fraction of aged MT lattices at the central spindle might explain why a model of uniform amplification of all EGFP-α-tubulin-labelled lattices predicted an excess of MT plus-ends in pole-distal regions (Fig. S3 D). To test this, we simulated MT plus-end distributions using the measured SiR-tubulin fluorescence profiles as a template (Fig. S3 I). Indeed, these simulations yielded a MT plus-end density along the spindle axis that fit the measured MT plus-end distributions more accurately, with the standard error of the estimate decreasing by 43%. This suggests that the frequency distribution of acentrosomal nucleation events approximates that of SiR-tubulin. Hence, a model of uniform amplification of SiR-tubulin-stained MT lattices can better explain the observed distribution of MT plus-ends.

Next, we tested whether this model can also explain the altered MT plus-end distributions in Augmin-depleted cells (Fig. S2 C-F). To visualize long-lived MT lattices, we treated control and HAUS6-depleted cells with SiR-tubulin (Fig. 3 F). EB3-EGFP imaging verified that SiR-tubulin treatment did not affect the distribution of MT plus-ends (Fig. 3 F, G). HAUS6 depletion led to a reduction in the integrated mass of SiR-tubulin-stained MTs at distances from the pole greater than 2 μm (Fig. 3 H), showing that Augmin depletion affects the distribution of aged MT lattices in the spindle. Thus, for each condition, we predicted MT plus-end distributions using the measured SiR-tubulin distributions as templates. We found that the model predicted the observed MT plus-end distribution in control cells (Fig. 3 I), as well as in HAUS6-depleted cells, but only when we accounted for the reduction in the available pool of amplification factors (Fig. 3 J, Fig. S3 J). Thus, our model can explain the spatial distribution of MT plus-ends observed in control metaphase spindles and predict the consequences of depleting Augmin. Furthermore, the prediction for MT plus-end distribution in HAUS6 RNAi cells was less accurate when the SiR-tubulin distribution of control cells was used as template (Fig. S3 J, blue dotted line), despite adjusting for the reduced size of the Augmin pool. Therefore, the model is sensitive to changes in the spatial distribution of template MTs.

Our data is consistent with a model in which Augmin binds to and amplifies aged MT lattices, as its preferential localization to cold-stable MTs would suggest. An alternative explanation is that Augmin distributes evenly across all spindle MTs but has increased nucleation activity near the poles. To distinguish between these possibilities, we investigated the spatial distribution of Augmin in fixed metaphase spindles by co-staining them with antibodies targeting HAUS6 and α-tubulin. We found that, like SiR-tubulin (Fig. 3, C-E) and acetylated-tubulin (Fig. S3, G and H), HAUS6 was enriched in pole-proximal spindle regions, relative to α-tubulin (Fig. S3, K and L). Combined, our modelling and immunofluorescence results suggest that Augmin gives rise to the MT plus-end distribution in metaphase cells by preferentially localizing to and thus amplifying aged MT lattices.

### MT growth direction is highly biased towards individual kinetochores

Unlike centrosomes, which generate point-symmetric radial arrays of MTs, Augmin generates MTs that grow at shallow angles relative to the parental MT (Kamasaki et al., 2013; Petry et al., 2013). Since many parental MTs in the mitotic spindle are bound to kinetochores, this directional bias could promote efficient formation of kinetochore-fibers. Indeed, Augmin depletion has been shown to impair kinetochore-fiber assembly (Bucciarelli et al., 2009; Goshima et al., 2008; Uehara et al., 2009; Lawo et al., 2009; Zhu et al., 2008). However, MT branch points are relatively scarce in electron tomograms of mitotic cells (Kamasaki et al., 2013). *In vitro*, Augmin-mediated branching can generate bundles of parallel MTs, but the presence of TPX2, a spindle associated factor that preferentially localizes to kinetochore-fibers (Bird and Hyman, 2008), stimulates the formation of splayed MT arrays instead (Petry et al., 2013). Thus, whether Augmin-mediated branching generates MTs with focused growth towards individual kinetochores has remained unclear.

To test whether MT growth is biased towards kinetochores during kinetochore-fiber assembly, we visualized individual growing MT plus-ends and kinetochores in early prometaphase spindles imaged by lattice light-sheet microscopy (Fig. 4 A, Movies S5 and S6). Using the poles as fiduciaries, we constructed regions of interest for each time point of the movie, to determine whether and when MTs begin to grow preferentially towards the ensemble of chromosomes. We specified two regions around each spindle pole, either facing the ensemble of kinetochores or facing outwards to the cell cortex (Fig. 4, B and C) and profiled the number of MT plus-ends in each region as a function of distance from the pole (Fig. 4, D-G). Immediately after nuclear envelope disassembly, neither region showed an increase in the number of MT plus-ends with distance from spindle pole. However, as early as 30 s after nuclear envelope disassembly, the number of MT plus-ends increased significantly at pole-distal regions facing kinetochores, but not in regions facing outwards (Fig. 4 D). The number of MT plus-ends growing towards the kinetochore ensemble continued to increase throughout mitosis, such that within 10 minutes after nuclear envelope disassembly their spatial distribution within the spindle was indistinguishable from that observed in steady-state metaphase spindles (Fig. 4 F). Thus, MT growth increases in spindle regions facing towards the ensemble of chromosomes during early stages of spindle assembly.

**Figure 4.**
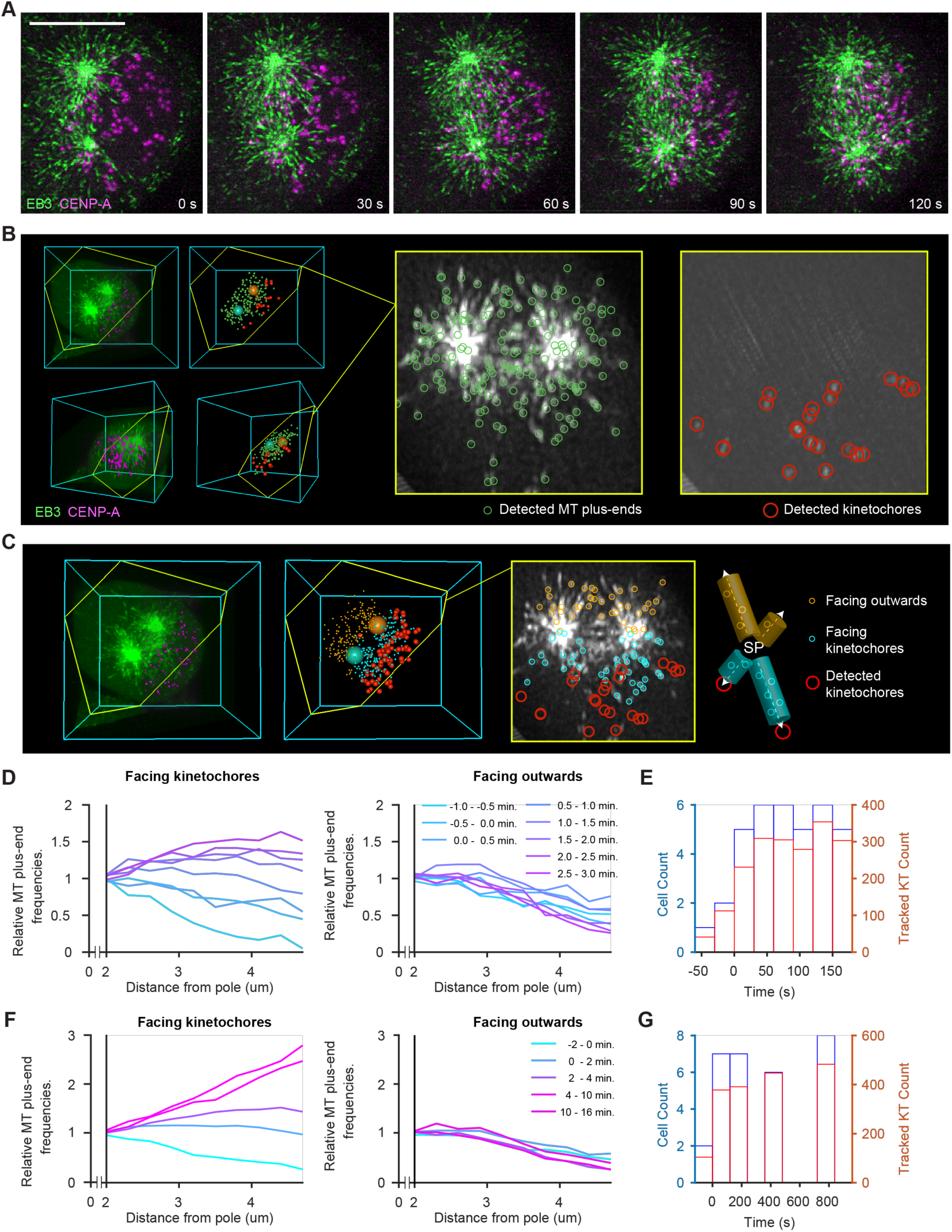
Non-centrosomal MTs increase in spindle regions facing towards kinetochore already during early prometaphase. (A-C) HeLa cell expressing EB3-EGFP (green) and mCherry-CENPA (magenta), imaged during prometaphase by 3D lattice light-sheet microscopy. (A) Maximum-intensity projections of deconvolved movie frames (t = 0 s, nuclear envelope disassembly). (B) Automated 3D detection of MT plus-ends and kinetochores in EB3-EGFP and CENP-A images, respectively. The slice highlighted in yellow follows the plane defined by the spindle poles and a randomly chosen kinetochore. The thickness of the projection is 600 nm. (C) Distribution of plus-ends mapped to regions “facing kinetochores” and “facing outwards”. MT plus-ends “facing kinetochores” are mapped in 500 nm wide cylinders around individual pole-to-kinetochore axes whereas MT plus-ends “facing outward” are mapped to “mirror” cylinders facing the opposite direction. (D) Quantification of detected MT plus-ends at increasing distances from the nearest spindle pole, throughout early prometaphase (t < 5 min, n > 268 movie frames analyzed per minute). Counts in individual cells are normalized to the value at 2 μm distance (n = 9 cells). (E) Histograms of cell and kinetochore counts for data shown in D. (F) Quantification of MT plus-ends from early prometaphase to metaphase. Counts in individual cells are normalized to the value at 2 μm distance from the pole (n = 22). (G) Histograms of cell and kinetochore counts for data shown in (F). Scale bars, 10 μm.

It has been proposed that molecular gradients generated around chromatin stabilize MTs growing towards chromosomes during early spindle assembly (Athale et al., 2008). This could at least partially explain the higher MT plus-end densities measured in Fig. 4, but is not expected to cause a bias towards individual kinetochores, relative to adjacent regions. To elucidate the MT growth direction relative to individual kinetochores, we first inspected z-sections in which isolated kinetochores were in focus with a spindle pole. In early prometaphase cells, we observed several instances in which many MT plus-ends grew along the respective pole-to-kinetochore axis, at much higher density than in adjacent regions (Fig. 5, A and B). This was more prominent in metaphase cells, in which many MT plus-ends grew along trajectories leading to kinetochores (Fig. 5, C and D). These observations support a model where a MT bound to a kinetochore promotes directional growth of additional MTs along its lattice.

**Figure 5.**
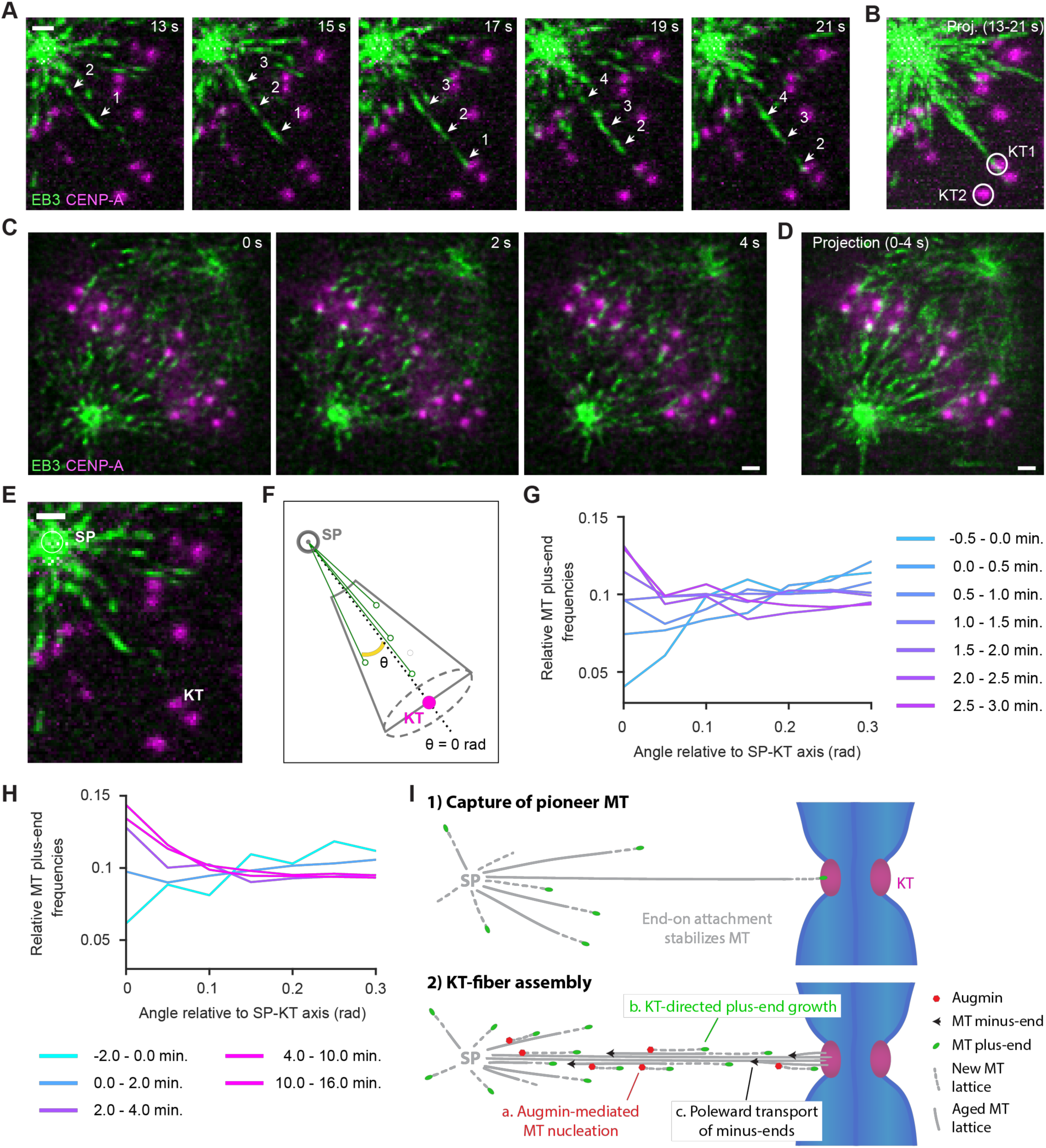
MT growth direction is highly biased towards kinetochores. (A-E) Lattice light-sheet microscopy of EB3-EGFP and mCherry-CENPA-expressing HeLa cells. (A-B) Kinetochore-directed MT growth in early prometaphase cell; detail of movie shown in Fig. 4 A (t = 0 s, nuclear envelope disassembly). (A) Each frame is a maximum-intensity projection of 5 z-sections, the center section being the focal place of the kinetochore highlighted in (B, KT1, white circle). A series of MT plus-ends are shown growing towards the kinetochore (arrow heads). (B) Maximum-intensity projection of movie shown in (A). (C) Kinetochore-directed MT growth in a metaphase cell, imaged at one stack of 50 z-sections/s. Each frame shows a single z-section. (D) Temporal projection of 4 s interval, as shown in (C). (E, F) Illustration of assay for automated quantification of MT growth direction relative to pole-kinetochore axes. (E) Spindle poles and kinetochores were automatically detected as in Fig. 4 to calculate conical region-of-interests centered on pole-kinetochore axes. MT plus-ends at distance < 2 μm to the pole were not considered, as they were not reliably resolved as individual objects. (F) MT plus-ends were mapped in conical regions-of-interest and for each, the angle of an axis connecting to the spindle pole was calculated relative to the pole-kinetochore axis. Solid green circles on black arrows illustrate radial positions of MT plus-ends. (G and H) Radial distributions of MT plus-ends relative to spindle-pole-kinetochore axes. Each curve represents data from all detected kinetochores and MT plus-ends for the indicated time interval relative to nuclear envelope disassembly (0 min). (I) A model for Augmin-driven assembly of kinetochore (KT)-fibers. Once “pioneer” MTs generated at the spindle poles (SP) attach to KTs, they become templates for Augmin-mediated amplification. Augmin localizes preferentially to aged MT lattices and generates an increasing fraction of spindle MTs (a.). This leads to a measurable KT-directed bias in MT plus-end growth (b.). Together with stabilization and poleward transport of MTs (c.), this amounts to a strong positive feed-back loop conducive to rapid KT-fiber assembly. Scale bars, 1 μm.

To quantify the directional bias in MT growth, we established a computational procedure to map the distribution of MT plus-ends relative to individual kinetochores. Based on the automatically detected positions of spindle poles and kinetochores, we defined cone-shaped 3D regions-of-interest connecting spindle poles with each individual kinetochore, beginning at a minimal distance of 2 μm from the spindle pole (Example illustrated in Fig. 5 E, F). We then measured the angle between the pole-kinetochore axis and each pole-MT plus-end axis to profile MT plus-end densities according to different angles relative to the pole-kinetochore axis (Fig. 5 F). To analyze the full movies, we tracked the pole and respective kinetochore over time and integrated MT plus-end positions of all kinetochores and all time points. Before and immediately after nuclear envelope disassembly, MTs did not preferentially grow towards kinetochores (Fig 5 G, -0.5 to 1.5 minutes). Thereafter, however, MT plus-end density increased substantially along directions pointing towards kinetochores (Fig. 5 G, H). Hence, the cytoplasmic space is not equally explored by growing MTs, but highly biased towards kinetochores as the spindle matures.

### Conclusions

Our analysis of MT plus-end distribution reveals the relative contributions of centrosomal and acentrosomal MT generation to mitotic spindle assembly in human cells. Although centrosome nucleation generates most MTs during the initial moments of spindle assembly, the fraction of MTs generated within the main spindle body increases rapidly after nuclear envelope disassembly. By metaphase, about 90% of the MTs arriving at chromosomes are generated in pole-distal regions, which compensates for the dilution of MT density in the radial arrays generated by centrosomes. This compensation was previously shown to enable the assembly of very large MT asters in *X. laevis* egg extracts (Ishihara et al., 2014; 2016). Our work shows that non-centrosomal MT nucleation is the predominant source of spindle MTs also in human somatic cells. Importantly, our analysis of HAUS6-depleted cells shows that the Augmin complex is responsible for this activity.

Mathematical modelling indicates that Augmin-mediated amplification of preexisting MTs largely explains how cells generate MT plus-end densities in interpolar regions of the metaphase spindle. The observed distribution of MT plus-ends is best explained by a model of uniform amplification of aged MT lattices, which we found to be selectively labelled by the live-cell dye SiR-tubulin. This suggests that the association time of Augmin with MT lattices may be rate-limiting for nucleation of new MTs. *In vitro* experiments with purified components indeed showed that a fraction of Augmin associates with MTs for ~3-13 s (Hsia et al., 2014). This behavior might explain why, also *in vitro*, <3% of branching points are found near the plus-end of the parent MT (Petry et al., 2013). While we can’t exclude that Augmin can mediate *de novo* nucleation on the lattices of short-lived MTs, our data suggests it preferentially associates with and amplifies long-lived MTs. The increasing abundance of non-centrosomal MTs during progression from prometaphase to metaphase could therefore reflect the increasing number of long-lived, kinetochore-bound MTs.

If the majority of spindle MTs are nucleated away from centrosomes, the establishment of robust pole-to-kinetochore attachments requires that their minus-ends move efficiently towards the poles. Indeed, prior work showed that the minus-ends of acentrosomal MTs are transported polewards by motor proteins (Lecland and Lüders, 2014). This transport slows down in pole-proximal regions, resulting in a local accumulation of MT minus-ends – an observation consistent with our own finding that aged MT lattices are enriched in these regions. It is known that the formation of the large *X. laevis* spindle relies on the sorting of MTs nucleated away from the poles (Yang et al., 2007; 2008; Brugués et al., 2012; Decker et al., 2018; Needleman et al., 2010). This study highlights the importance of further characterizing motor activity in human somatic spindles. Our mathematical model explains how amplification of a steady-state MT template network gives rise to the distribution of growing MT plus-ends during metaphase; future efforts must be aimed at understanding how the stabilization and motor-dependent reorganization of nascent MTs gives rise to the evolving template network.

Lattice light sheet microscopy revealed a directional bias of MT growth towards individual kinetochores. This might be explained by the shallow angle of Augmin-nucleated MTs relative to the parental MT (Kamasaki et al., 2013; Petry et al., 2013). However, kinetochore-directed MT growth also often occurred along curved trajectories, particularly in metaphase spindles, suggesting that the respective plus-ends remain associated with MTs that have already attached to kinetochores. This might be mediated by interactions between MT lattices, as observed in mature kinetochore-fibers (Petry et al., 2013; Nixon et al., 2015), or by plus-end-associated factors forming compliant connections with neighboring MT lattices (Molodtsov et al., 2016).

Overall, our study shows that kinetochore-fibers assemble by coordinated growth of multiple MTs, rather than by a sequence of independent stochastic search-and-capture events. Our data support the existence of a positive feedback mechanism, whereby the capture of pioneer MTs during early prometaphase generates a long-lived template for Augmin-driven amplification (Fig. 5 I). Having the vast majority of spindle MTs nucleated on and guided along previously captured MTs imposes a strong directional bias which, coupled to poleward transport of minus-ends, is conducive to efficient bundle formation. As metaphase spindles continue to turn over their MTs, kinetochore-directed MT growth might further contribute to maintenance of kinetochore-MT interactions and the resolution of incorrect merotelic attachments. Our quantitative assays provide a resource to further study this and the mechanisms underlying kinetochore-directed MT guidance.

## Acknowledgments

The authors thank Kai Johnsson and Luc Reymond for providing SiR-tubulin and SiR-Hoechst, Grazvydas Lukinavicius and Iva Tolic for comments on the manuscript, IMBA/IMP/GMI BioOptics for technical support, and Life Science Editors for editing assistance. Imaging data used in this publication was produced in collaboration with the Advanced Imaging Center, a facility jointly supported by the Gordon and Betty Moore Foundation and HHMI at HHMI’s Janelia Research Campus. Research in the laboratory of D. W.G. has been supported by the European Community’s FP7/2007-2013 under grant agreements nr. 241548 (MitoSys) and nr. 258068 (Systems Microscopy), by an ERC Starting Grant (nr. 281198), by the Wiener Wissenschafts-, Forschungs- und Technologiefonds (WWTF; project nr. LS14-009), and by the Austrian Science Fund (FWF special research program SFB “Chromosome Dynamics”; project nr. SFB F34-06). Research in the laboratory of G.D. has been supported by the NIH grant R01 GM067230 and the Human Frontier Science Program (fellowship LT000954/2015 to P.R.).

## Author contributions

Conceptualization: DWG, AD. Formal analysis: AD, PR. Funding acquisition: DWG, EB, GD. Investigation: AD, PR. Methodology: AD, PR, WL. Project administration and supervision: DWG, EB, GD. Visualization: AD, PR. Writing: DWG, AD, with input and approval by all authors.

## Materials and Methods

### Cell lines and cell culture

HeLa cell lines stably expressing fluorescent reporter proteins were derived from a HeLa Kyoto line obtained from S. Narumiya (Kyoto University, Japan) and validated by a Multiplex human Cell line Authentication test (MCA). The EB3-EGFP/mCherry-CENPA and EGFP-α-tubulin/H2B-mCherry lines were previously published (Cuylen et al., 2016; Dick and Gerlich, 2013) and generated as described in (Schmitz and Gerlich, 2009). Briefly, reporter constructs were subcloned into IRESpuro2b or IRESneo3 vectors that allow expression of resistance genes and tagged proteins from a single transcript. The resulting plasmids were transfected into the parental cell lines using X-tremeGENE9 DNA transfection Reagent (Sigma-Aldrich) according to the manufacturer’s instructions. The EB3-tagRFP line was generated by transfecting a plasmid derived from the EB3-eGFP construct published in (Stepanova et al., 2003): the eGFP sequence was replaced with tagRFP (from Evrogen Cat.# FP142) by restriction enzyme cloning. For selection of reporter construct expression, cells were cultured in medium containing 500 μg/ml G418 (Thermo Fisher, Cat. No. 11811-031) and 0.5 μg/ml puromycin (Merck Millipore, Cat. No. 540411).

The RPE1 cell line stably expressing EB3-EGFP and mCherry-CENPA was generated from a parental hTERT-RPE1 line obtained from ATCC, using a lentiviral vector system previously described (Samwer et al., 2017). For selection of reporter construct expression, cells were cultured in 10 μg/mL blasticidin (Sigma-Aldrich, Cat. No. 15205).

HeLa and RPE1 cells were cultured in Dulbecco’s modified Eagle medium (DMEM; produced in-house at IMBA, Austria) supplemented with 10% (v/v) fetal bovine serum (FBS; Thermo Fisher), 1% (v/v) penicillin-streptomycin (Sigma-Aldrich) and GlutaMAX (Thermo Fisher), at 37°C with 5% CO_2_ in a humidified incubator. All cell lines used in this study have been regularly tested negatively for mycoplasma contamination.

### Plasmids

We aimed to use minimally tagged HAUS6 variants. To monitor expression levels via EGFP, we designed plasmids containing the P2A sequence (2A peptide from porcine teschovirus-1 polyprotein, (Szymczak et al., 2004)). The ribosome fails to insert a peptide bond at the two last amino acids of the P2A sequence, yielding two separate polypeptides from a single mRNA. To generate EGFP-P2A-HAUS6, a HAUS6 cDNA clone was obtained from the PlasmID Repository (pCR-BluntII-TOPO_HAUS6; see Key Resources Table for details) and amplified by PCR with the following primers: 5’- aagagaatcctggacc gaccggtatgagctcggcctcg - 3’ (forward) and 5’- ctggat cggaattcggatcctcatcttgtcaagtcagacg -3’ (reverse). Alternatively, to generate the EGFP-P2A control construct, amplification with the forward primer 5’- aagagaatcctggaccgaccggtatgagctcggcctcggtcacc -3’ was used to introduce a stop codon two amino-acids downstream of the start methionine of HAUS6. The amplified sequences were inserted via Gibson Assembly (New England Biolabs) into a previously described EGFP-P2A-BAF_IRES_Blast construct (Samwer et al., 2017), replacing the BAF gene. To generate an siRNA-resistant HAUS6 variant, we introduced five mismatches in the siRNA target region without changing the amino-acid sequence (see Fig. S2 H for sequence details). The mutated gene region was produced by gene synthesis (gBlocks^®^ by Integrated DNA Technologies) and swapped into the EGFP-P2A-HAUS6 plasmid described above by Gibson Assembly. All plasmids were verified by DNA sequencing and will be distributed via Addgene.org. For the RNAi phenotype complementation experiments, the EGFP-P2A-HAUS6* and EGFP-P2A -encoding plasmids were delivered to EB3-tagRFP - expressing cells 24h after siRNA transfection (see below). In each reaction, 200 ng of plasmid were transfected using X-tremeGENE9 DNA transfection Reagent (Roche) according to the manufacturer’s instructions.

### SiRNA transfection

SiRNAs (see Key Resources Table) were delivered at a final concentration of 20 nM, using Lipofectamine RNAiMAX (Thermo Fisher). 6.8 pmol siRNA were dissolved in 20 μl OptiMEM; 2 μl RNAiMAX were diluted in 20 μl OptiMEM. Both solutions were combined, mixed by pipetting and incubated for 20-30 min. Cells were harvested by trypsinization, resuspended in fresh DMEM medium and seeded onto LabTek II chambered coverglass (Thermo Fisher). 2 x 10^4^ HeLa cells or 3 x 10^4^ RPE1 cells were seeded per well, in 300 μL DMEM medium. 40 μl of the above transfection mix were added dropwise to the cells directly after seeding. Cells were analyzed 48 h later by confocal live-cell microscopy (or lysed for quantification of protein levels). Alternatively, cells were harvested by trypsinization 24 h after siRNA transfection and then seeded onto pre-cleaned 5 mm coverslips, for analysis by Lattice light-sheet microscopy 24 h later.

### Immunoblotting

Where quantification of protein levels by immunoblotting was preformed, samples were prepared in parallel as those meant for live-cell imaging. All steps were done at RT. HeLa cells treated as above were lysed in 1x SDS loading buffer 48 h after siRNA transfection. Samples were separated by Novex NuPAGE SDS-PAGE system (Thermo Fisher) using 4-12% BisTris 1.5 mm gels in MES running buffer, according to manufacturer’s instructions. Proteins were transferred to a nitrocellulose membrane (Protran BA 83, Sigma-Aldrich) by semidry blotting in a Trans-Blot^®^ SD Cell (BioRad). Membranes were blocked in 5% (w/v) milk powder + 0.02% NP-40 in TBS (blocking solution) for 30 min, then incubated for 1.5 h with the primary antibodies diluted in blocking solution at the concentrations indicated in the Key Resource Table. Membranes were washed three times in blocking solution (sequentially for 5, 10, and 10 min), then incubated for 1.5 h with species-specific secondary antibodies coupled to horseradish peroxidase (HRP) at the concentrations indicated in the Key Resource Table. Membranes were washed three times in blocking solution for 5 min, then three more times in 0.02% NP-40 in TBS (sequentially for 5, 10, and 10 min). Finally, membranes were rinsed once with ddH_2_O and incubated for 5 min in Pierce™ ECL Plus Western Blotting Substrate (Thermo Fisher). Chemiluminescence was documented on a ChemiDoc™ MP (BioRad) system. All immunoblots were recorded with no saturated pixels.

### Immunofluorescence

HeLa Kyoto WT cells were cultured as described above. All subsequent steps were done at room temperature, unless otherwise stated. See Key Resource Table for antibody dilutions used. Cells were fixed for 6 minutes in -20° C methanol, then washed three times with PBS supplemented with Tween 80 (PBST), each for 5 min. Samples were blocked with 10% fetal calf serum in PBST (blocking solution) for 30 min and then incubated with the primary antibody in blocking solution for 14 h, at 4 C°. Samples were washed two times with PBST, each for 10 min, then incubated with the respective secondary antibody in blocking solution for 3 h. Samples were washed for 10 min in PBS (repeated three times) and then imaged on a spinning-disk confocal microscope (UltraView VoX, PelkinElmer) controlled by Volocity software, with a 100x/1.45 NA oil objective. Two 3D volumes were sequentially acquired by illuminating with 488 nm and 561 nm lasers.

### Confocal live-cell microscopy

Cells were imaged on LabTek II chambered coverglass (Thermo Fisher) in DMEM containing 10% (v/v) FBS and 1% (v/v) penicillin-streptomycin, but without phenol red and riboflavin to reduce autofluorescence (Samwer et al., 2017). Where indicated, the imaging medium additionally contained 50 nM SiR-Hoechst or 50-100 nM SiR-tubulin (as specified in the respective figure legends), and cells were pre-incubated in it for >2 h. Cells were maintained at 37 °C in a humidified atmosphere of 5% CO_2_, provided by incubation chambers (European Molecular Biology Laboratory (EMBL), Heidelberg, Germany) installed on every microscope used.

To study the spatial distributions of EB3-EGFP-labeled MT plus-ends, α-tubulin-labeled MTs, and SiR-tubulin-stained MTs, fast time-lapse imaging was performed on a spinning-disk confocal microscope (UltraView VoX, PelkinElmer) controlled by Volocity software, with a 100x/1.45 NA oil objective. Single z-slices were acquired at 1-2 s/frame, for a total of 30 s to 1 min (as indicated in the respective figure legends).

To investigate the timing of SiR-tubulin binding to growing MT lattices, HeLa cells stably expressing EGFP-α-tubulin were imaged on an LSM880, AxioObserver microscope equipped with an Airyscan detector and controlled by ZEN 2011 software, with a Plan-Apochromat 63x/1.4 NA oil objective (Zeiss). Single z-sections were acquired at 1 s/frame, for a total of 2 minutes, in Super-resolution mode.

### Lattice light-sheet microscopy

Cells were grown on pre-cleaned 5 mm coverslips and maintained in DMEM containing 10% (v/v) FBS and 1% (v/v) penicillin-streptomycin, but without phenol red at 37 °C. Lattice light sheet microscopy was performed on the instrument described in (Chen et al., 2014). Briefly, the coverslips were mounted on the microscope in CO_2_-independent L15 medium containing 10% FBS, without phenol red and maintained at 37 °C for the duration of the experiment.

In experiments where unperturbed cells were imaged (Fig. 1, 4 and 5), we first identified prophase cells based on increased EB3-EGFP density around asters; the exclusion of soluble EB3-EGFP from nuclear areas indicated that the nuclear envelope was still intact. Using widefield time-lapse microscopy, we monitored mitotic progression for every cell. We recorded the onset of prometaphase, defined by the influx of cytoplasmic EB3-EGFP into the nucleus due to nuclear envelope disassembly. We then switched to fast 3D lattice light-sheet imaging, performed using a 15 μm long square excitation lattice pattern of outer numerical aperture equal to 0.50 and inner numerical aperture equal to 0.42. For each cell, 3D two color volumes of EB3-EGFP and mCherry-CENPA were acquired by sequentially illuminating each plane with 488 nm and 560 nm lasers. 75-150 time points were acquired at a volumetric imaging rate of 1 Hz. After imaging, 44 out of 45 cells subsequently entered anaphase, indicating minimal phototoxicity. For analysis of HAUS6-depleted spindles (Fig. 2), we identified metaphase cells based on spindle morphology and kinetochore congression to the equatorial plane. 3D lattice light-sheet imaging was performed using an excitation pattern of outer numerical aperture equal to 0.55 and inner numerical aperture equal to 0.44. For each cell, 3D two color volumes of EB3-EGFP and mCherry-CENPA were acquired by sequentially illuminating each plane with 488 nm and 589 nm lasers. 150 time points were acquired at a volumetric imaging rate of 1 Hz. All images were acquired in sample-scan imaging mode with a lateral translation of 0.4 μm and subsequently deskewed in post processing. Final voxel dimensions for all lattice light-sheet image datasets were 104 nm x 104 nm x 210-217 nm.

### Processing and analysis of confocal microscopy images

#### Spindle Registration

Wherever confocal time-lapse movies were recorded for analysis of the average spatial distribution of fluorescent markers in the metaphase spindle, the movie frames were first registered using the MultiStackReg Fiji plugin (Thévenaz et al., 1998) to correct for translation and rotation of the spindle (rigid body transformation). Registration of EB3-EGFP movies was first performed on filtered images (Gaussian Blurr: σ = 5 pixel) to make sure individual plus-ends did not impact the transformation. If EB3-EGFP movies were collected, registration was performed on them and the transformation matrices obtained subsequently applied to unfiltered images of all channels. Where EB3 was not imaged (Fig. 3 C-E, S3 A), registration was based on the EGFP-α-tubulin instead.

#### Fluorescence intensity profiles from the spindle pole

To study the spatial distribution of fluorescent markers in spindles imaged by live-cell microscopy, we quantified the respective fluorescence intensities in average temporal projections of the registered time-lapse movies. Only cells that had both poles in the same focal plane were imaged. To study the distribution of acetylated tubulin and HAUS6 in immunofluorescence images, we quantified fluorescence intensities in the central plane of suitably oriented spindles, with both poles in focus.

We analyzed the distribution of fluorescence intensity in a polar coordinate system centered around one of the spindle poles (*d* = 0). The reference axis is a manually drawn line extending towards the second pole. The fluorescence intensity was measured along a series of circumferential line profiles (radius increment of 1 px = 130 nm) placed between the first spindle pole and the cell equator. The mean and total intensities measured within the inner ⅓ (interpolar) section of the profiles were normalized to the cytoplasm of individual cells and then to the mean value measured at d = 520 nm (or 4 px), taken as the rim of the centrosome.

#### Fraction of MT plus-ends not attributable to the pole

The number of MT plus-ends in imaged spindles could not be directly measured by detection of individual EB3-EGFP spots, as detection accuracy was limited by spot density and unreliable in pole-proximal regions of the metaphase spindle. Therefore, we inferred the relative number of MT plus-ends from the mean EB3-EGFP fluorescence intensity *I*, measured as a function of distance from pole (*d*) as described above. The measured curves should reflect the distribution of plus-end density *D* along the spindle axis:

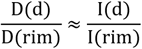

The spherical geometry of centrosomal MT growth predicts that MT plus-end density decreases by the inverse of the squared distance from the origin (*d*= *rim*). Thus, the relative number of plus-ends *N* is given by

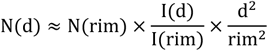

The probability that a dynamically instable MT nucleated at the centrosome grows to a certain length depends on the catastrophe and growth rescue rates, which are in turn thought to be length-dependent (Foethke et al., 2009; Wordeman and Stumpff, 2009). If this were the case, the number of MT plus-ends growing from centrosomes is expected to decrease exponentially as a function of distance from pole. If the number of growing MT plus-ends at the centrosome rim is *N(rim)*, the number expected at any given distance *d* is

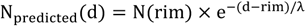

The characteristic MT length λ is itself a function of the parameters describing the dynamic instability behavior, namely the frequencies of catastrophe (*f_cat_*) and rescue (*f_res_*) over time:

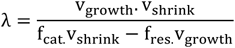

where v_growth_ and v_shrink_ are the growth and shrinkage rates of dynamic MTs (Verde et al., 1992). The decay constant *k* that governs the distribution of plus-end number is 1 / *λ*. We calculated a decay constant k of 0.08 μm^-1^ based on catastrophe and rescue frequencies measured in the outer spindle regions of LLCPK-1α cells (Rusan et al., 2001). At any given distance *d*, the fraction of plus-ends that cannot be explained by this model of dynamic instability is thus:

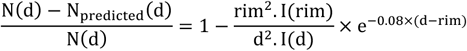

#### EB3-EGFP and SiR-Hoechst intensities from the metaphase plate

The fluorescence intensities were measured in average temporal projections of registered time-lapse movies and normalized to the cytoplasm of individual cells. Mean intensity profiles were taken along a 10-pixel-thick line connecting the center of the spindle (i.e. the midpoint of the interpolar axis) to one of the poles. Measurements were then taken along a line of the same length and thickness, also parallel to the interpolar axis, displaced sideways to the spindle periphery. Both profiles were normalized to the mean of the first 5 values measured inside the spindle body.

#### EB3-EGFP intensity along pole-KT trajectories

To minimize distortions caused by kinetochore motion, we quantified EB3-EGFP fluorescence in projections of 6-second intervals of our registered movies. Maximum-intensity projections were generated for 3 sequential frames using the Running ZProjector Fiji plugin. In selected frames of the projections, 5-pixel-wide segmented lines were drawn, connecting selected kinetochore pairs to one of the spindle poles (Fig. 1 I). To avoid measurement artifacts resulting from the periodic kinetochore oscillations, CENPA loci that showed little motion blur were selected; the drawn lines were typically curved, following the natural curvature of kinetochore-fibers. The fluorescence intensities measured along the lines to the cytoplasm of individual cells were normalized in all three imaged channels. All profiles were aligned to the mid-point between the two sister kinetochores, identified as local maxima in the mCherry-CENPA profile. Fluorescence intensities were normalized to the average values measured in a 5-pixel-wide window around the last local maxima, i.e. the kinetochore closest to the spindle pole.

#### Model of MT template amplification

At any given distance from the spindle pole (*d*), the total number of MT plus-ends attributable to centrosomal activity is predicted to be N(rim) x e^-k(d-rim)^, where *rim* is the estimated radius of the centrosome. Due to the spherical geometry of centrosomal MT arrays, a decreasing fraction of these plus-ends is visible in the thin optical planes we analyze:

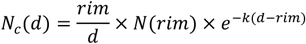

Our model adds to this array of centrosome-generated plus-ends those expected to result from MT branching, i.e. *de novo* generation on the lattices of preexisting MTs, N_b_:

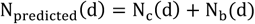

While the component *N_c_(d)* is solely a function of the number of MT plus-ends generated at the spindle poles (and directly inferred from the mean EB3-EGFP fluorescence measured at *d = rim*), *N_b_(d)* will depend on the frequency of MT plus-end generation across the entire spindle. It has been proposed that the activity of MT nucleation factors can be spatially modulated by chromatin-generated molecular gradients. Notably, the RanGTP effector TPX2 has been shown to stimulate Augmin-mediated MT branching *in vitro* (Petry et al., 2013). By interacting with preexisting MTs, Ran-activated factors like TPX2 can reach high concentrations in the main body of human mitotic spindles (Oh et al., 2016). However, it remains unclear whether TPX2 activity varies significantly within the spindle body – and how acentrosomal MT nucleation rates may vary as a result. The concentration gradients of both RanGTP and TPX2 appear rather diffuse, reaching all the way from chromosomes to the spindle poles (Kalab et al., 2006; Oh et al., 2016). In summary, there is currently no evidence of a differential in acentrosomal nucleation activity along the axis of human mitotic spindles. Hence, for simplicity, we assumed in our model that the MT branching frequency *Bra* is not spatially modulated, not depending on distance from the pole but only on the local abundance of branching factors.

Since Augmin’s association with MT minus-ends is only transient, lost shortly after the nucleation event and as the poleward transport begins, we further assume that the branching factors do not accumulate at pole-proximal regions of the spindle, continuously redistributing across template MTs to drive local generation of MT plus-ends. The local branching activity *Bra(d)* is then a linear function of the template distribution *Template(d)*, depending only on the total amount of branching factors available (*Aug*):

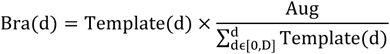

It is crucial to consider not just *Bra(d)* – the plus-ends generated at *d* in any given instant of time – but also the plus-ends that were generated in preceding instants (at all distances from the pole *a, a ∈] rim, d]*) and grew to reach *d* in the intervening time. Since Augmin is known to nucleate MTs at shallow angles, with the same polarity as the parent lattices (Kamasaki et al., 2013; Petry et al., 2013), we assumed that plus-ends generated by branching share the directionality of the template network, their angle to the centrosome remaining constant. Further, we assumed that their growth is governed by the same dynamic instability parameters as centrosomal MTs. The probability that a MT generated by branching at *d* = *a* reaches any given distance from pole is thus p(d) = e^−kx(d−a)^, where *k* is the decay constant that depends on catastrophe and rescue rates. The total number or plus-ends generated by branching is given by

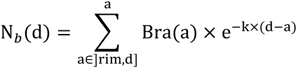

The two components combine to generate a total number of plus-ends given by

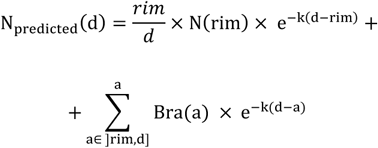

We infer *Template(d)* from the observed distribution of either EGFP-α-tubulin or SiR-tubulin fluorescence, quantified in live cells as described above. Since we don’t know how the measured fluorescence units relate to MT number, both *Template(d)* and *Bra(d)* are unitless. Consequently, the amount of branching factors *Aug* is also an abstract quantity – and the one free parameter in the model. In every simulation, we predicted plus-end distributions for a range of *Aug* values (0-5 in 0.1 increments) and chose the one that minimized the mean standard error of the estimate MSE (see Fig. S3 D for an example):

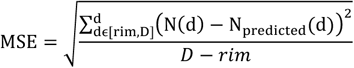

A Kolmogorov-Smirnov test was used to compare the best-fitting predicted distribution with the total EB3-EGFP fluorescence measured in cells. The resulting statistics are given in the respective figure legends.

This mathematical model was implemented in custom written MATLAB code, made available with this publication.

#### Quantification of SiR-tubulin binding to MT lattices in interphase cells

Instances where an EGFP-α-tubulin -labeled lattice grew into a region of low MT density were identified by visual inspection. Mean EGFP-α-tubulin and SiR-tubulin fluorescence intensities were then measured in rectangular ROIs placed in front of the growth event (each 350 nm in width and 1.4-1.6 μm in length, as depicted in Fig. 3 A), in each of the subsequent 60 frames. The background fluorescence values measured at t = 0 s (when the lattice has not yet grown into the ROI) was subtracted from each of the two channels, respectively. The profiles were finally normalized to the mean value measured at t > 30 s.

### Analysis of lattice light-sheet microscopy data

The analysis of lattice light-sheet microscopy data was performed with the U-track 3D framework (Roudot et al, in preparation) developed in Matlab (Mathworks). Of a dataset of 65 cells imaged by lattice-light sheet microscopy, 39 contained the entire spindle volume and had sufficiently homogeneous signal-to-background ratio for automated MT plus-end detection. These were considered for further analysis. Each movie represented 1-2 min intervals, and together covered mitotic stages from 2 min before nuclear envelope disassembly until 16 minutes after nuclear envelope disassembly. The automatic analysis pipeline consists of three steps: i) definition of dynamic region-of-interest inside the spindle (dROI), ii) MT plus-end detection or intensity sampling iii) measurement mapping in dROI to interrogate MT plus-end statistics and integration across non-synchronous movies.

#### Dynamical ROIs definition

The high concentration of MT plus-ends at the spindle pole results in a bright signal that moves with the whole spindle. As such, these large clusters are excellent fiducials to build a dROI for the spindle and associated frame of reference.

The spindle poles are detected in a statistical framework that first establishes a set of candidate objects using the size of detected clusters in the 3D volume. A scale map is estimated using 3D Laplacian of Gaussian filtering at multiple scales (Lindeberg, 1998). The scales range from twice the size of a diffraction limited spot (see Section “MT plus-end detection”) to 2 μm, using steps of 100 nm. The candidate locations are the local maximum across all filtered scales. Those candidate locations are then tracked over time using a Brownian motion prior with a maximal inter-frame displacement of 1 μm and allowing temporary disappearance of up to one frame (Jaqaman et al., 2008). Each resulting track is then scored according to the product of its lifetime and median intensity. The two best candidates are selected as the spindle poles.

Kinetochore are also tracked over time in order to interrogate MT plus-end location statistics between the spindle poles and moving kinetochores. Each kinetochore’s location is estimated using a 3D implementation of the algorithm described in (Aguet et al., 2013). Sharp transitions in kinetochore motion between pre- and post-MT capture make trajectory estimation difficult with conventional methods. To solve this problem, we applied a recent algorithm for tracking erratic motion via piecewise-stationary motion modeling to each kinetochore trajectory (Roudot et al., 2017). The combination of pole and kinetochore trajectories enables the creation of multiple dROIs and associated frame of reference for quality control in 3D data and statistical analysis.

#### MT plus-end detection

MT plus-ends are detected using the centroid of the region masked with the adaptive thresholding algorithm described in (Aguet et al., 2013). This algorithm requires a single parameter, an estimate of the scale of the diffraction-limited objects. In order to estimate the scale of a diffraction limited MT plus-end, the scale of each object is approximated using the fitting of a 3D Gaussian function and the set of resulting scales is fitted with a Gaussian mixture model. The mode of the resulting distribution is kept as the estimated scale. Quality control was performed by automatic rendering of dROI and detection overlay in Matlab. All MT plus-ends were counted, regardless of their growth direction.

#### 3D measurements of fluorescence intensity in LLSM movies

Intensity measurement inside the spindle are carried out through random sampling inside the 3D region of interest with a constant density of 100 loci/μm^3^.

#### Quantification of MT plus-end densities facing kinetochores and outward

The definition of regions of interest facing kinetochore” and “facing outward” to study MT plus-end density at different distances from spindle poles was performed using a collection of cylindrical ROIs between the spindle poles and each kinetochore with a radius of 500 nm. Each kinetochore is associated to only one pole (the closest pole at the end of the trajectory), resulting in two sets of dROIs 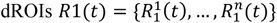 and 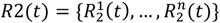. Outward-facing areas are defined through point reflection of each dROI using the spindle pole as a center. Let us denote P the set of labeled plus-end and ***x***\* the coordinate of plus-end in the frame of reference associated to its closest dROI 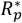. The histogram of distance from spindle pole of detected MT plus-end is defined as:

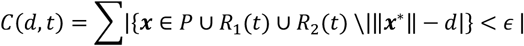

Where |.| denotes the set cardinality, d is the polar distance ranging from 2 to 6 μm, and *ϵ* is a binning parameter set to 150 nm. In order to make measurement comparable and interpretable, relative frequencies counts rather than probability distributions are shown:

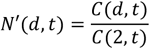

Integrating count statistics of multiple ROI also has to take into account the different length of each ROI. To do so, each relative count according to the number of dROI sampling the respective distance was taken into account.

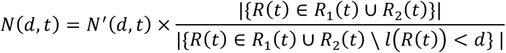

Where *l*(.) denotes the length of the dROI. Quality control for this normalization process was performed on multiple movies presenting stationary metaphase. The same process was performed symmetrically for outward-facing dROIs.

#### Quantification of MT growth direction relative to pole-kinetochore axis

To determine the density of MTs growing along the pole-kinetochore axis relative to adjacent regions, kinetochores were analyzed individually as illustrated in Fig. 5, using conical dynamical ROIs with an angle of 0.5 radian. Let us denote 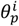 the elevation of a detected MT plus-end in the dROI 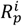, the angle histogram was defined as

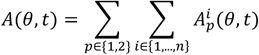

with

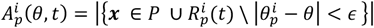

Where *θ* is the polar distance ranging from 0 to 0.5 radian and *ϵ* is a binning parameter set to 0.025 radian. As MT plus-ends can be counted multiple times and the changes in dROI shape over time must be taken into account considering the volume of a 3D cone, let us denote angular histograms normalized as

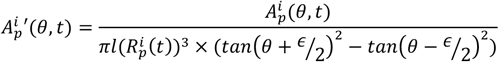

This normalization enables to integrate thousands of dROI across space, time and acquisitions. The probability distributions, *A*(*θ*, *t*), were then computed for different time intervals relative to nuclear envelope disassembly as shown in Fig. 5.

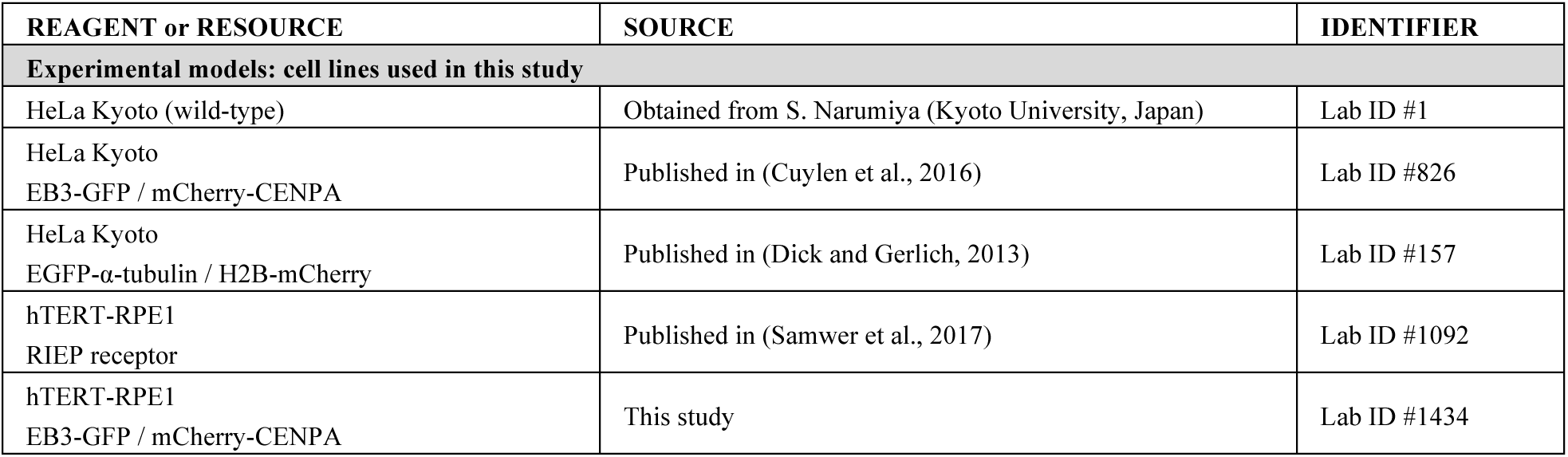
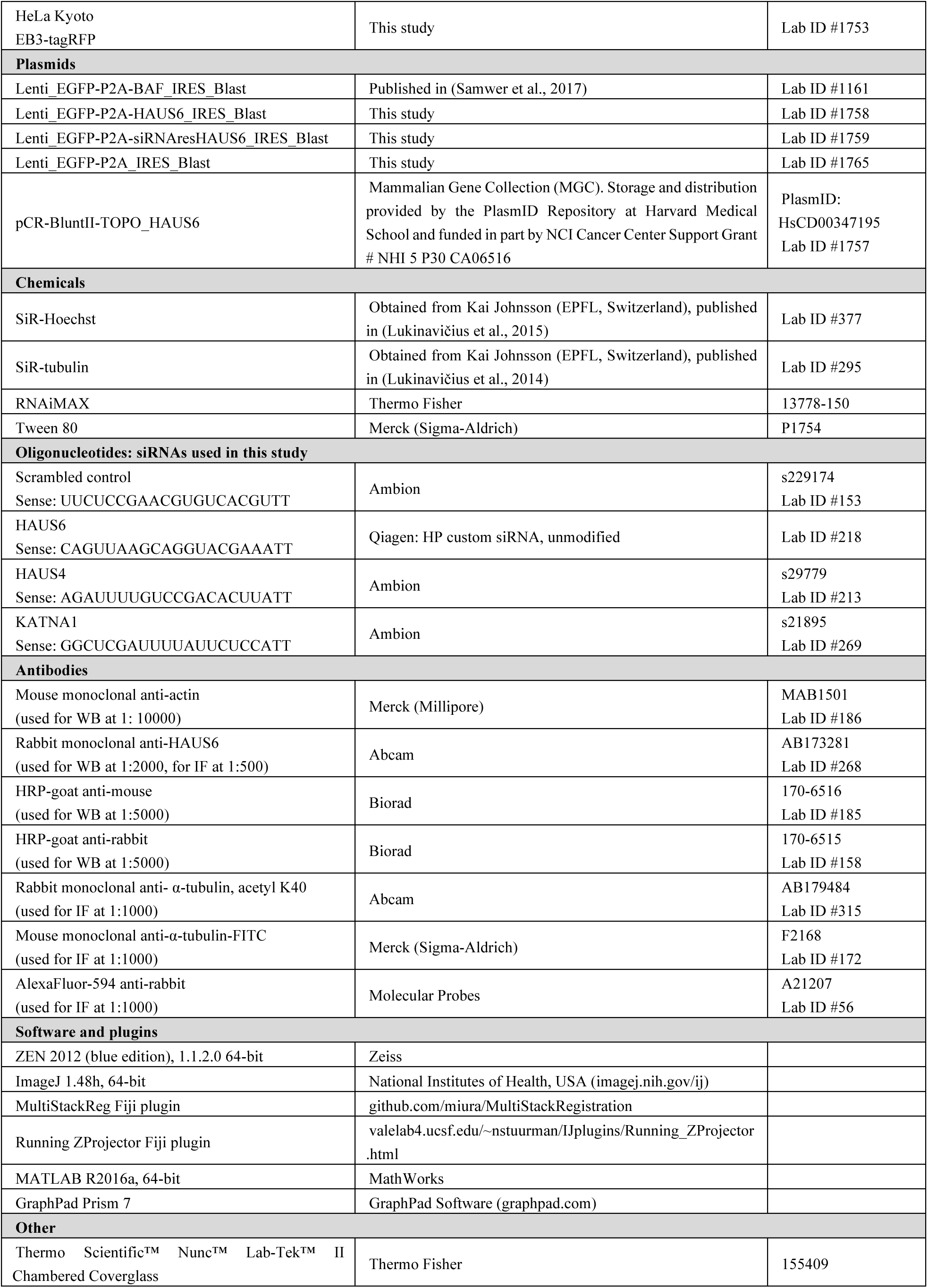
Key resources table.

---

**Supplementary figure 1.**
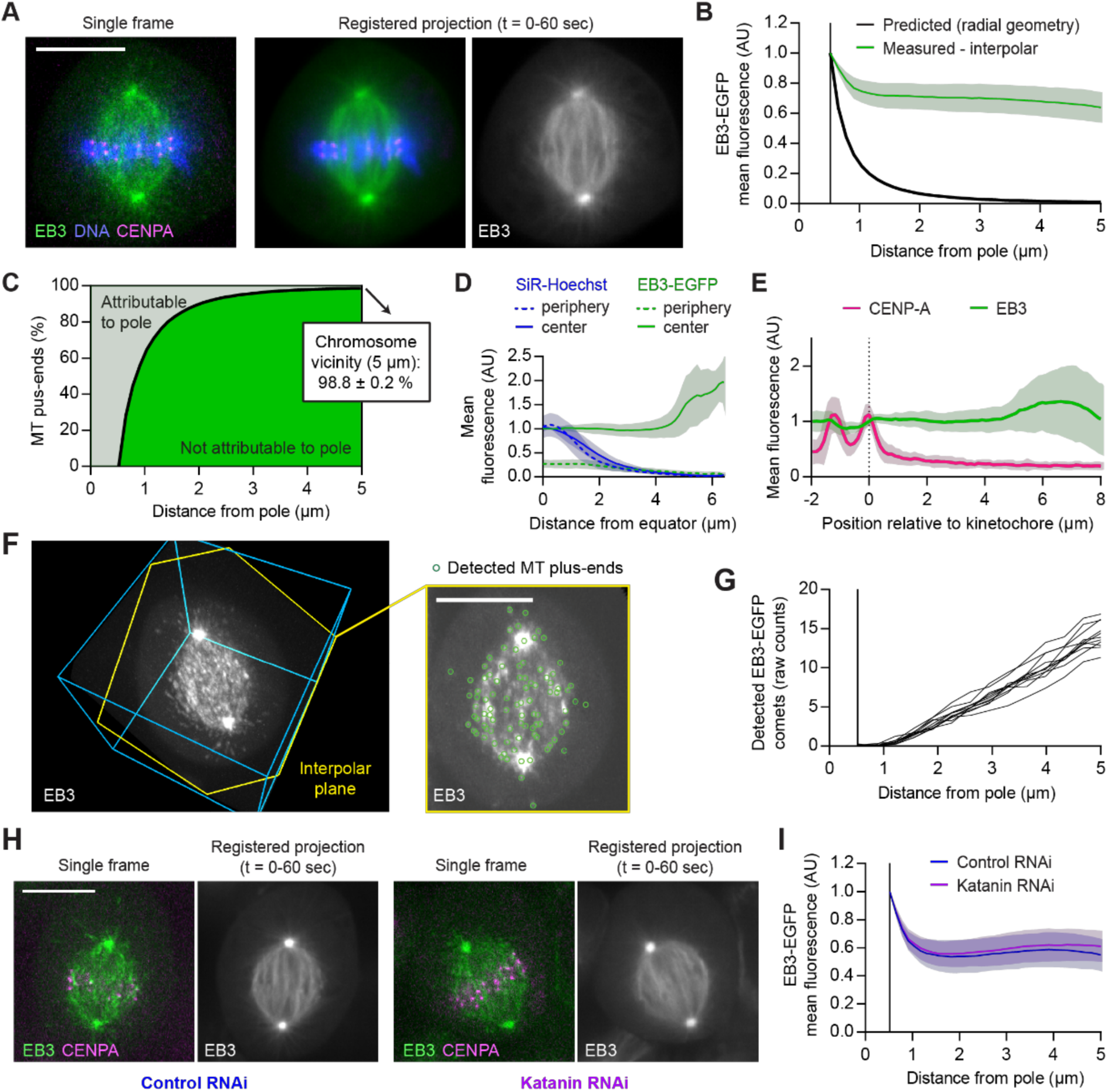
(A-B, D-E) Live-cell confocal microscopy of hTERT-RPE1 cells expressing EB3-EGFP (green) and mCherry-CENPA (magenta), stained with SiR-Hoechst (blue) (n = 25 cells). (A) 1-minute time-lapse movies were acquired during metaphase at 2 s/frame (see Movie S2), registered to correct for spindle rotation (see Materials and Methods for details) and projected to obtain mean-intensity images for analysis of MT plus-end distributions. (B) Mean EB3-EGFP fluorescence intensity quantified in the projections thus obtained as in Fig. 1 B-C. Black line indicates predicted signal dilution by radial geometry. Individual measurements were normalized to centrosome rim. (C) Fractions of MT plus-ends attributable (light-green) and not attributable (dark-green) to nucleation at the spindle poles, in HeLa metaphase spindles. Computed from the measurements shown in Fig. 1 C as detailed in Materials and Methods. (D) Mean EB3-EGFP and SiR-Hoechst fluorescence measured across the chromosome-cytoplasm boundary as described in Fig. 1 D. (E) Mean EB3-EGFP and mCherry-CENPA fluorescence along curved lines connecting pairs of sister-kinetochores to one of the spindle poles. Quantified as described in Fig. 1 F-G (n = 49 profiles in 8 cells). (F) Automated 3D detection of MT plus-ends in EB3-EGFP images. The slice highlighted in yellow follows the plane defined by the spindle poles and a random kinetochore. (G) Quantification of detected MT plus-ends at increasing distances from the nearest spindle pole. Plotted are the mean plus-end numbers per movie frame, for each cell (n=11). (H and I) EB3-EGFP and mCherry-CENPA-expressing HeLa cells transfected with control or Katanin-targeting siRNAs (n= 14 and 18 cells, respectively). (H) Cells imaged during metaphase by live-cell confocal microscopy, for 1 min at 1 s/frame. (I) Mean EB3-EGFP fluorescence intensity quantified in average-intensity projections of registered time-lapse movies, as described in Fig. 1 B-C.

**Supplementary figure 2.**
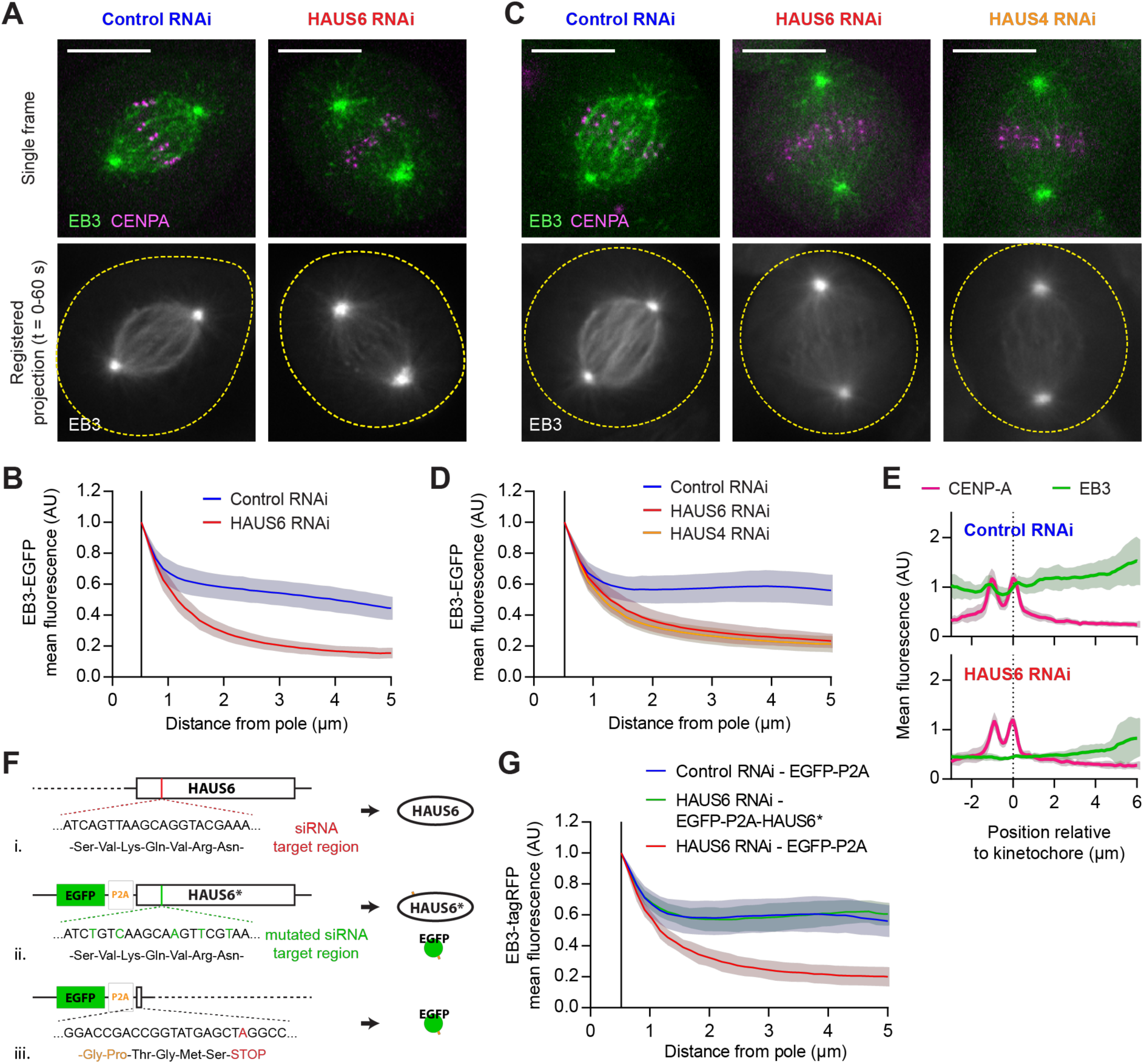
(A-E) Live-cell confocal microscopy of EB3-EGFP and mCherry-CENPA-expressing cells. (A and B) hTERT-RPE1 cells transfected with control or HAUS6-targeting siRNAs (n = 34 and 31 cells, respectively, collected in 3 independent experiments). (C-E) HeLa cells transfected with control, HAUS4-targeting or HAUS6-targeting siRNAs (n = 33, 34 and 36 cells, respectively, collected in 3 independent experiments). (A and C) 1-minute confocal time-lapse movies were acquired during metaphase at 2 s/frame. Dashed yellow lines indicate cell boundaries. (B and D) Mean EB3-EGFP fluorescence intensity quantified in average-intensity projections of registered time-lapse movies, as described in Fig. 1 B-C. (E) Mean EB3-EGFP and mCherry-CENPA fluorescence along curved lines connecting pairs of sister-kinetochores to one of the spindle poles. Quantified as described in Fig. 1 F-G (n = 29 and 22 profiles for control and HAUS6 RNAi cells, respectively; in 7 cells per condition). (F and G) EB3-tagRFP -expressing cells were transfected with control or HAUS6-targeting siRNAs. (F) Plasmids encoding for EGFP-P2A-HAUS6* (F.ii, for expression of siRNA-resistant HAUS6) or EGFP-P2A (F.iii, control) were delivered 24h later. See Materials and Methods for details. (G) 48h after siRNA transfection, cells were imaged during metaphase as in C and E. Mean EB3-tagRFP fluorescence was measured in EGFP positive cells as described in Fig. 1 B-C. Lines and shaded areas denote mean ± s.d., respectively. Scale bars, 10 μm.

**Supplementary figure 3.**
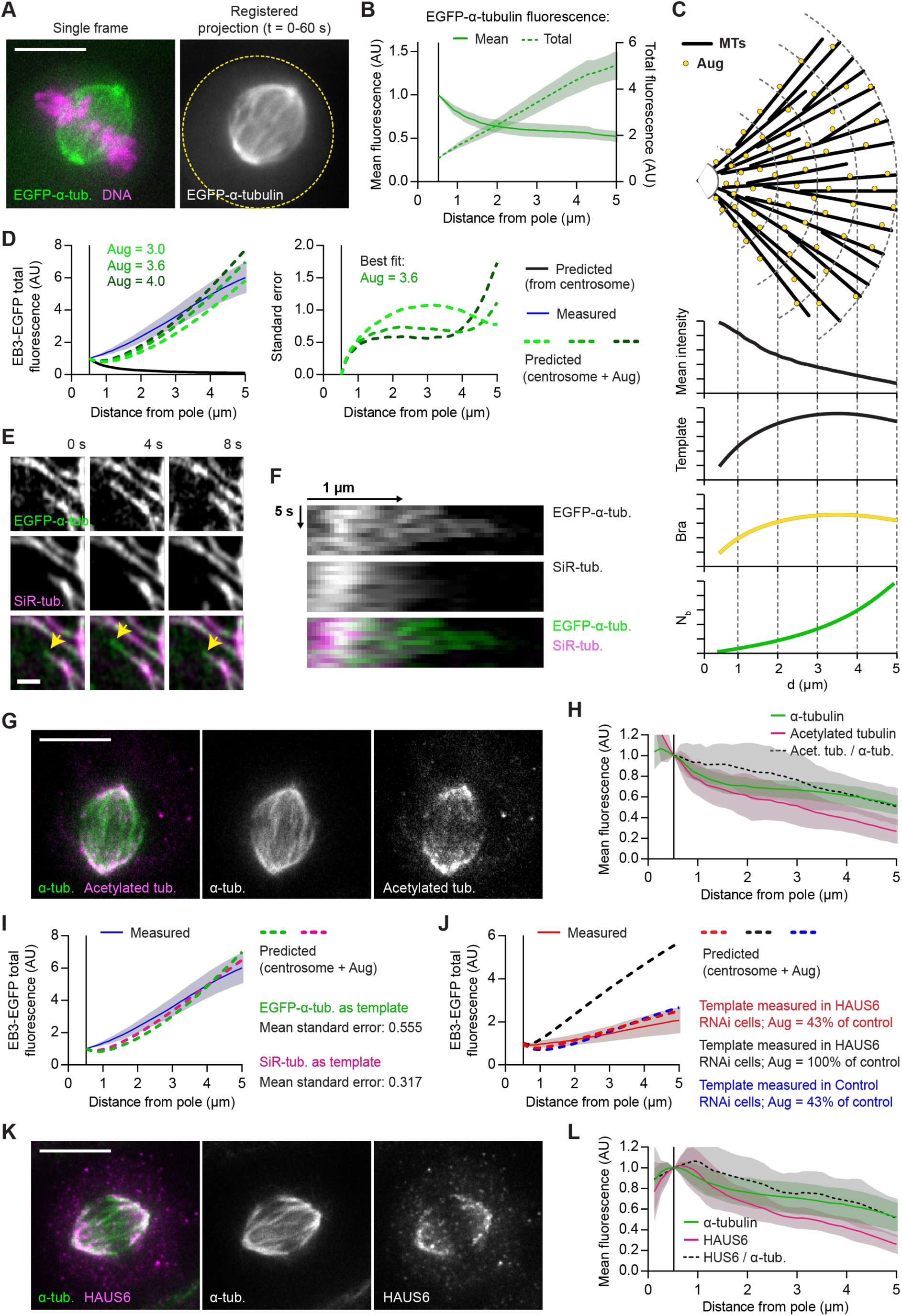
(A and B) Live-cell microscopy of HeLa cells expressing EGFP-α-tubulin (green) and H2b-mCherry (magenta). (A) 1-minute time-lapse movies were acquired during metaphase at 2 s/frame, registered to correct for spindle rotation and projected to obtain mean-intensity images for analysis of MT lattice distributions. Dashed yellow line denotes cell boundary. (B) Mean and total EGFP-α-tubulin fluorescence quantified in interpolar spindle regions as in Fig. 1 B. (C) Schematic representation of a model for uniform amplification of a template MT network. MT branching factors (Aug) distribute across MT lattices and generate plus-ends at a fixed rate. The frequency distribution of MT-dependent plus-end generation (Bra) is thus set to be a linear function of template distribution (Template), which is inferred from the mean fluorescence intensity of a MT marker (measured as in A-B). The number of MT plus-ends generated on MT lattices (Nb) is a function of the branching frequency distribution Bra. For a prediction of MT plus-end distributions, Nb is added to the number of plus-ends originating at the spindle poles. See Materials and Methods for details. (D) Three separate mathematical simulations wherein the template network measured in B is amplified using the indicated amplification parameters (Aug). The predicted MT plus-end distribution is compared to the total EB3-EGFP fluorescence measured in Fig. 1 A-C (left). The standard error of each prediction is plotted against distance from pole (right). (E-F) Interphase HeLa cells expressing EGFP-α-tubulin and incubated with 100 nM SiR-tubulin were imaged at 1 s/frame. (E) Example of a growing MT undergoing a catastrophe event (arrow heads point to EGFP-labeled growing tip). (F) Kymogram of the growth event shown in E. (G-H) Acetylated tubulin and α-tubulin visualized by immunofluorescence in metaphase spindles of HeLa cells. (H) Mean fluorescence intensities quantified in interpolar regions as in Fig. 1 B-C (n = 26 cells collected in two independent experiments). (I and J) Mathematical simulations of MT template amplification. The predicted totals of MT plus-ends are compared to the total EB3-EGFP fluorescence profiles measured (I) in Fig. 1 A-C or (J) in Fig. 3 J. (I) The measured fluorescence of the indicated MT markers (plotted in Fig. 3 E) was used to estimate MT template distributions. (J) The SiR-tubulin fluorescence measured in Fig. 3 H (in control or HAUS6 RNAi cells, as indicated) was used to estimate MT template distributions. (K-L) HAUS6 and α-tubulin visualized by immunofluorescence in metaphase spindles of HeLa cells. (L) Mean fluorescence intensities quantified in interpolar regions as in Fig. 1 B-C (n = 34 cells collected in two independent experiments). Lines and shaded areas denote mean ± s.d., respectively. Scale bars, 10 μm.

